# Direct and indirect impacts of positive selection on genomic variation in *Drosophila serrata*

**DOI:** 10.1101/2022.03.31.486660

**Authors:** Yiguan Wang, Adam J. Reddiex, Scott L. Allen, Stephen F. Chenoweth

**Affiliations:** School of Biological Sciences, The University of Queensland, St Lucia, Australia; Biological Data Science Institute, The Australian National University, Australia; Institute of Evolutionary Biology, University of Edinburgh, Edinburgh EH9 3FL, UK

**Keywords:** *Drosophila serrata*, positive selection, selective sweep, machine learning

## Abstract

Understanding the extent to which microevolutionary adaptation relies on novel beneficial mutations, as opposed to previously neutral standing genetic variation, is an important goal of evolutionary genetics. Progress towards this goal has been enhanced during the genomic era through the study of selective sweeps. Selective sweeps fall into two categories: hard sweeps via new mutations and soft sweeps via pre-existing mutations. However, data are currently lacking on the relative frequency of these two types of selective sweep. In this study, we examined 110 whole genome sequences from *Drosophila serrata* sampled from eastern Australia and searched for hard and soft sweeps using a deep learning algorithm (diploS/HIC). Analyses revealed that approximately 15% of the *D. serrata* genome was directly impacted by soft sweeps, and that 46% of the genome was indirectly influenced via linkage to these soft sweeps. In contrast, hard sweep signatures were very rare, only accounting for 0.1% of the scanned genome. Gene ontology enrichment analysis further supported our confidence in the accuracy of sweep detection as several traits expected to be under frequent selection due to evolutionary arms races (e.g. immunity and sperm competition) were detected. Within soft sweep regions and those flanking them, there was an over-representation of SNPs with predicted deleterious effects, suggesting positive selection drags deleterious variants to higher frequency due to their linkage with beneficial loci. This study provides insight into the direct and indirect contributions of positive selection in shaping genomic variation in natural populations.

## Introduction

Evolutionary genetics views the process of adaptation from two broad perspectives. The quantitative genetic view focusses on phenotypic traits that respond to directional selection in proportion to their additive genetic variance (Falconer, 1996; Lush, 2013; Lynch & Walsh, 1998). Here, traits are assumed to be polygenic and so only subtle allele frequency changes are expected at underlying loci. Molecular population genetics historically has focused on the fate of individual mutations in response to positive selection, relying on predictable patterns of diversity changes (Barton, 1998; Smith & Haigh, 1974). A key distinction between these two views has been the degree to which adaptation is thought to be limited by the availability of beneficial mutations. Under the quantitative genetic view, most beneficial variants already exist as standing variation before the onset of selection contributing to a sometimes-rapid response. In contrast, under the classic molecular population genetic view, adaptation is limited by the rate at which beneficial mutations arise. More recently, the gap between these two quite different perspectives has begun to close through theoretical developments in the study of selective sweeps and new methods to detect them with population genomic data.

Positive selection can leave a signature on the DNA polymorphisms around the target site of selection through hitchhiking and recombination (Durrett & Schweinsberg, 2004; Kaplan et al., 1989; Smith & Haigh, 1974). For example, when a new advantageous mutation appears and survives stochastic loss from genetic drift, it will rapidly spread in the population and increase in frequency. The genomic region immediately surrounding the advantageous mutation will have reduced genetic diversity, a skewed site frequency spectrum and elevated levels of linkage disequilibrium (Nielsen, 2005; Sinha et al., 2011). The genomic signature left by such an episode is defined as a hard selective sweep (Hermisson & Pennings, 2005).

Selective sweeps can also arise from standing genetic variation when a pre-existing neutral or mildly deleterious allele becomes beneficial due to an environmental shift and change in selection (Hermisson & Pennings, 2005; Messer & Petrov, 2013). For example, the reduction of body armour plates in three-spined sticklebacks was likely caused by selection on standing genetic variation when they migrated from a marine environment into freshwater lakes (Colosimo et al., 2005). In contrast to hard sweeps, soft sweeps start from standing genetic variation and not mutation, therefore, the newly beneficial variant exists at a higher frequency and on multiple haplotypes. Because of this, soft sweeps do not skew the allele frequencies of linked neutral polymorphisms to the same extent as hard sweeps and the local reduction in genetic variation is also less extreme (Kern & Schrider, 2016; Przeworski et al., 2005). Specific examples of soft sweeps include lactase persistence alleles in humans (Jones et al., 2013), resistance to treatment in pathogens such as bacteria (Croucher et al., 2014), and plant height in wheat (Raquin et al., 2008). Soft sweeps may also play key roles in evolutionary rescue, a process that allows a declining population to recover from the demographic consequences of harsh environmental shifts (Wilson et al., 2017).

Different types of sweeps can result in different evolutionary dynamics and distinct genetic outcomes (Barrett & Schluter, 2008). For instance, alleles of small effect are more likely to contribute to adaptation through soft sweeps compared to hard sweeps, this is because the higher starting allele frequency of standing variants relative to new mutations leads to a higher probability of fixation of the beneficial allele. Additonally, soft sweeps may lead to more rapid adaptation as populations are not limited by the rate of new mutations, but have almost immediate access to beneficial variants. Soft sweeps can also have indirect population genetic consequences such as increasing the frequency of linked deleterious alleles (Hartfield & Otto, 2011). It is therefore clear that significant insights into the genetic process of adaptation at single loci can be gained by distinguishing soft from hard sweeps in natural populations.

Distinguishing soft from hard sweeps has proven challenging (Jensen, 2014). For example, a hard sweep signature can also arise from a soft sweep if the beneficial allele is present at a low enough frequency before the sweep begins. Similarly, a hard sweep from a novel beneficial mutation can appear soft if the sweep is ancient and recombination has degraded linkage of surrounding neutral sites (Garud et al., 2015). Furthermore, fluctuations in population size can also blur the distinction between hard and soft sweeps. For example, a severe bottleneck can make a soft sweep resemble a hard sweep (Wilson et al., 2014). Another challenge comes from the ‘shoulder effect’ of hard sweeps, which could generate spurious signatures of soft and partial sweeps (Schrider et al., 2015).

Despite these challenges, noticeable advances in detecting the signature of selective sweeps and distinguishing soft sweeps from hard sweeps have been made as more powerful and robust tools became available. For example, H12 and H2/H1(Garud et al., 2015), are tests based on haplotype homozygosity, that can detect recent soft and hard sweeps while being robust to demographic changes. More recently, diploS/HIC, which uses a convolutional neural network to classify genomic regions into hard and soft sweeps, linked to a sweep, or neutral, was proposed and has been applied to *Anopheles gambiae* (Kern & Schrider, 2018) and green Anolis lizards (Bourgeois & Boissinot, 2019). The method has high detection and clsssification accuracy while being reasonably robust to demographic history misspecification.

*Drosophila serrata* is a vinegar fly distributed along the north eastern coastal areas of Australia and much of Papua New Guinea (Jenkins & Hoffmann, 2001). The species has been used as a quantitative genetic model for studying climatic adaptation (Frentiu et al., 2009), sexual selection (Hine et al., 2002), sexual dimorphism (Chenoweth & Blows, 2003) and evolutionary constraints (Gosden et al., 2018; Hine et al., 2004). Multiple artificial selection experiments and experimental evolution studies founded from wild-derived collections confirm that the species harbours abundant standing variation (Blows & Hoffmann, 1993; Gosden et al., 2018; Rundle et al., 2005). Despite a detailed quantitative genetic understanding of adaptation and evolutionary potential in the species, we know little about how selection manifests at the genomic level. While latitudinal cline studies suggest that spatially varying selection likely affects gene expression (Allen, Bonduriansky, et al., 2017), the degree to which within-population genomic variation is subject to hard and/or soft sweeps is unknown. In this study, our goal was to understand how positive selection directly and indirectly impacts genomic variation in *D. serrata*. To achieve this we applied the powerful machine learning method diploS/HIC (Kern & Schrider, 2018), to 110 genomes sampled from a single endemic population of *D. serrata* from Brisbane, Queensland, Australia (Reddiex et al., 2018).

## Materials and Methods

### Fly collection and sequencing data

We used the sequence data available from the *Drosophila serrata* genomic reference panel (DsGRP) (Reddiex et al., 2018). Briefly, *D. serrata* females were collected from a single endemic population at Bowman Park, Brisbane, Queensland, Australia (Latitude: - 27.45922, Longitude: 152.97768) in 2011. Lines were immediately isogenised via 20 generations of full-sib mating to preserve natural genetic variation and minimise adaptation to the laboratory. All 110 lines were sequenced to approximately 23.5X coverage on an Illumina Hi-Seq 2000 sequencing platform using 100 base-pair paired-end reads with 500 base-pair inserts (Reddiex et al., 2018).

### SNP genotyping and quality filtering

Reads were mapped to the *D. serrata* reference genome (version 1.0) (Allen, Delaney, et al., 2017) with BWA-mem (version v0.7.10) (Li, 2013) using default settings. Reads mapping to indels were realigned using GATK IndelReagligner (version3.2-2) (McKenna et al., 2010). We implemented SNP calling using the Joint Genotyper for Inbred Lines (JGIL), a probabilistic model tailored to call SNPs for large panels of inbred lines with high accuracy (Stone, 2012). We set the minimum read mapping quality score and the minimum JGIL genotype quality score to 20, which both correspond to a 99% probability of being correct. Sites failing to meet the genotype quality score were treated as missing data and masked with an “N”. We then removed non-biallelic sites, excluded sites with read depths outside a range of 5 to 60, and masked any heterozygous sites (set as “N”). Repetitive sequences, annotated by Allen, Delaney, et al. (2017) using RepeatMasker version 3.0 (Smit et al., 1996-2010), were also removed in the data filtering. Finally, we excluded sites that were genotyped for less than 80% of the DsGRP lines. A diagram outlining this filtering approach can be found in **Supplementary Figure S1**. The data filtering resulted in a total of 11,820,435 SNPs for analysis.

### An updated D. serrata genome

The reference genome of *D. serrata* was created using long-read sequencing technology and has a length of 198 Mbp and a contig N50 of 0.94 Mbp (Allen, Delaney, et al., 2017). We subsequently used Dovetail™ HiRise and Hi-C methods to scaffold those contigs and achieved a scaffold N50 of 30.3 Mb. The six largest scaffolds span 80% of the genome and reach near chromosome-arm level length except for 2L, which is spanned by two large scaffolds of 21Mb and 8.7 Mb. Because our analyses are genomic window-based approaches, we conducted all analyses on these six scaffolds.

### Inferring selective sweeps using diploS/HIC

We applied a powerful supervised machine learning approach, diploS/HIC (Kern & Schrider, 2018), to identify the signature of selective sweeps in the *D. serrata* population. diploS/HIC uses simulated training data to train a classifier. The diploS/HIC pipeline computes a vector of 15 different population genetic summary statistics for each of eleven 5k base-pair sub-windows, and feds this 15 x 11 matrix of information as input to a convolutional neural network (CNN). The CNN is trained to classify the central sub-window as one of five signatures of selection: a hard sweep, a soft sweep, a window linked to a hard sweep (linkedHard), a window linked to a soft sweep (linkedSoft), and a “neutral” window unlinked to a sweep (Kern & Schrider, 2018). The summary statistics in the 15-element feature vectors include three site frequency spectrum SFS estimators: *π* (Tajima, 1983), θ_*w*_ (Watterson, 1975), θ_*H*_, Fay and Wu’s H (Fay & Wu, 2000) and Tajima’s D (Tajima, 1989), three moments of the distribution of a multilocus ‘genotype distance’ score, *g_kl_*, computed between all pairs of individuals (variance, skew, and kurtosis), four different haplotype-like metrics: the number of multilocus genotypes and H_1_, H_12_ and H_2_/H_1_ (Garud et al., 2015), two measures of linkage disequilibrium, *Z_ns_* (Kelly, 1997) and the maximum value of ω (Innan & Kim, 2004), and maxFDA. Together, these summary statistics have been shown to harbour significant information on the presence of hard and soft sweeps (Kern & Schrider, 2018).

We used *discoal* (Kern & Schrider, 2016) to generate the training data sets. To date, we have no information on the population history of *D. serrata*, so we referred to parameters from *D. melanogaster* and *D. simulans* when we ran the simulations. A principal concern was to determine how robust our population genomic inferences were to different demographic scenarios. For this reason, we simulated different demographic scenarios with the intent to assess the impact of demographic misspecification on sweep classification. For all of the demographic histories simulated, we used an exponential distribution for the recombination rate with a mean of ρ = 5×10^-9^ and upper bound of 3ρ (Garud et al., 2015), a mutation rate, μ, that was distributed *~Uniform (1×10^-9^, 1.6×10^-8^)* (Haag-Liautard et al., 2007; Keightley et al., 2014), a selection coefficient, *s*, that was distributed *~Uniform (0.001,0.1)*, and initial frequency of selected loci for soft sweeps *Uniform(0, 0.2)*. Relatively recent selective sweeps were modelled by randomly setting the fixation time for each sweep between 0 and 160 generations ago. The simulation was performed on 55Kb length windows, each divided into 11 5Kb sub-windows. For simulated selection scenarios, a single locus in the centre of a sub-window was chosen as the target of selection. The target of selection was assigned to the central sub-window for sweep scenarios and one of the 10 non-central windows for linked selection scenarios. For the neutral scenario where no selected locus was assigned, we ran simulations with *s* set to zero. In total, this produced 46,000 simulations; 22,000 for hard sweeps, 22,000 for soft sweeps, and 2,000 for neutral scenarios.

We followed instructions from (https://github.com/kern-lab/diploSHIC/wiki/A-soup-to-nuts-example) to calculate the feature vectors of summary statistics on the simulations. The simulation data was then split into training (80%) and validation (20%) to optimise the number of epochs. Optimisation ran until the validation accuracy stopped improving beyond 0.001. This was reached after 15 epochs and a validation accuracy of 83.0%. To test the trained classifier and assess its accuracy on unseen data, predictions were made on an independent dataset simulated with the same parameter settings described above for the training data. Repetitive regions were masked when we converted *D. serrata* data into feature vectors. Using the trained CNN, *D. serrata* genomic windows were classified into hard sweeps, soft sweeps, linkedHard regions, linkedSoft regions and neutral regions.

The demographic history of the natural population from which the DsGRP was derived is currently unknown. However, a previous study on this population suggested that it might have experienced a bottleneck given the pervasive genome-wide negative Tajima’s D (Reddiex et al., 2018). We therefore sought to evaluate the sensitivity of our approach to this source of uncertainty. To evaluate the robustness of our results to different demographic histories, we trained three diploS/HIC classifiers based on simulation of the following scenarios: constant population size of 1,000,000, a short severe bottleneck where the population size contracted from 1,000,000 (1*N_0_*) to 33,000 (*0.033N_0_*) between time 0.330 and 0.340 (backward in units of *4N_0_*); and a long shallow bottleneck followed by slight population expansion where the population size was reduced from 1*N_0_* to 0.589*N_0_* at time 0.160 before recovering and expanding to 1.47*N_0_* by time 0.216. In addition to testing the prediction accuracy on independently simulated datasets with matching demographic histories, we also assessed the prediction accuracy from misspecified demographic history by, for example, using the classifier trained on constant population size simulations to make predictions on simulations from a bottleneck demographic history.

### Annotating the phenotypic impact of SNPs within genomic windows

Having classified the *D. serrata* genome into regions likely affected by different evolutionary processes, our next goal was to determine whether these regions differed in the number of functionally important mutations they contained. To achieve this, we first predicted the functional impact of SNPs in our datasets using SnpEff (Version 4.3t) (Cingolani et al., 2012) and then used SnpSift (Version 4.3t) (Ruden et al., 2012) to classify the SNPs into one of four degrees of functional impact: high, moderate, low and modifier. For example, variants that create a premature stop codon or remove a start codon are high impact variants whereas missense variants are considered moderate impact. Synonymous sites are considered low impact and intergenic variants are considered to be modifiers owing to their potential to affect transcription.

We used a permutation test to check if the SNP annotation features were enriched in any of the five genomic classes determined by diploS/HIC (Schrider & Kern, 2017). First, adjacent genomic windows receiving the same classification by diploS/HIC were merged into a single region. We then randomly assigned the same set of region lengths to the genome and counted the number of mutations belonging to the four SNP impact categories assigned by SnpEff/SnpSift. The data was then permuted 10,000 times. This test allowed us to check if the sweep regions were significantly enriched with SNPs predicted to have different functional impact severity while controlling for genomic region sizes. We also tested Gene Ontology (GO) enrichment using this permutation approach, but instead of counting predicted SNP effects, we counted GO terms of all genes annotated within each region. By taking this approach we were able to account for any inherent non-random distribution of GO terms across regions of the *D. serrata* genome.

### Alternate inferences of sweeps using H12 and piHS scan

In addition to diploS/HIC, we applied two commonly used haplotype-based methods, H12 (Garud et al., 2015) and *p*iHS (Gautier & Naves, 2011) to detect selective sweeps. H12 is based on haplotype homozygosity and was developed to detect recent selective sweeps (Garud et al., 2015). H12 calculates the frequencies of the first and second most common haplotypes within a predefined genomic window. It can be used to identify both hard and soft sweeps with similar power and is expected to be robust to various demographic scenarios (Garud et al., 2015). Moreover, soft and hard sweeps can be distinguished using an additional statistic H2/H1, where H2 is haplotype homozygosity calculated using all but the most frequent haplotype in a predefined window, and H1 is the haplotype homozygosity calculated using all haplotypes (Garud et al., 2015). For a selective sweep, the higher the H2/H1 value, the softer the selective sweep is. In the scan, we used the default parameter settings to perform the H12 scan: window size 400 SNPs, jump 50 SNPs, haplotype distance threshold 0, and H12 threshold 0.022.

*p*iHS scan (Gautier & Naves, 2011) is derived from iHS (Voight et al., 2006), a measure of whether a SNP is on an unusually long haplotype carrying the ancestral or derived allele. For iHS, a negative score indicates long haplotypes around the derived focal allele, while a positive score indicates the long haplotype around the ancestral focal allele (Urbinati et al., 2016). Since both high positive and high negative iHS scores are of interest, we usually transform iHS score into *p*iHS using the following equation which allows positive values and negative values on the same scale:

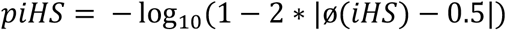

where, ø(x) represents the Gaussian cumulative distribution function (Gautier & Naves, 2011). The R package “rehh” (version 2.0.2) (Gautier & Vitalis, 2012) was used to perform *p*iHS scan for each polymorphic site. An FDR-adjusted *P*-value was used as the threshold, which was equivalent to *p*iHS=4.08.

## Results

### Performance of diploS/HIC

We first used simulations to evaluate the performance of diploS/HIC in classifying different types of selective events. In broad terms, the method performed very well under all three demographic scenarios (**Figure 1a**; **Supplementary File S1**) and misspecification of demographic history had very little impact on prediction accuracy (**Figure 1b**). For this reason, we focus on the constant population size scenario for the majority of the results. Based on a constant population size, it was extremely rare for a hard sweep or linkedHard region to be misclassified as a soft sweep, linkedSoft, or neutral, and vice-versa. The classifier was most accurate for soft sweeps, linkedSoft, and neutral. Prediction accuracy on the independent simulations for soft sweeps, linkedSoft, and neutral regions was very high (mostly >90%). Accuracy for hard sweeps was 72% and linkedHard had a median accuracy of 68%. Accuracy for linkedHard regions ranged from 44%-78% with misclassification of linkedHard as a hard sweep becoming more likely the closer a sub-window got to the central window.

**Figure 1.**
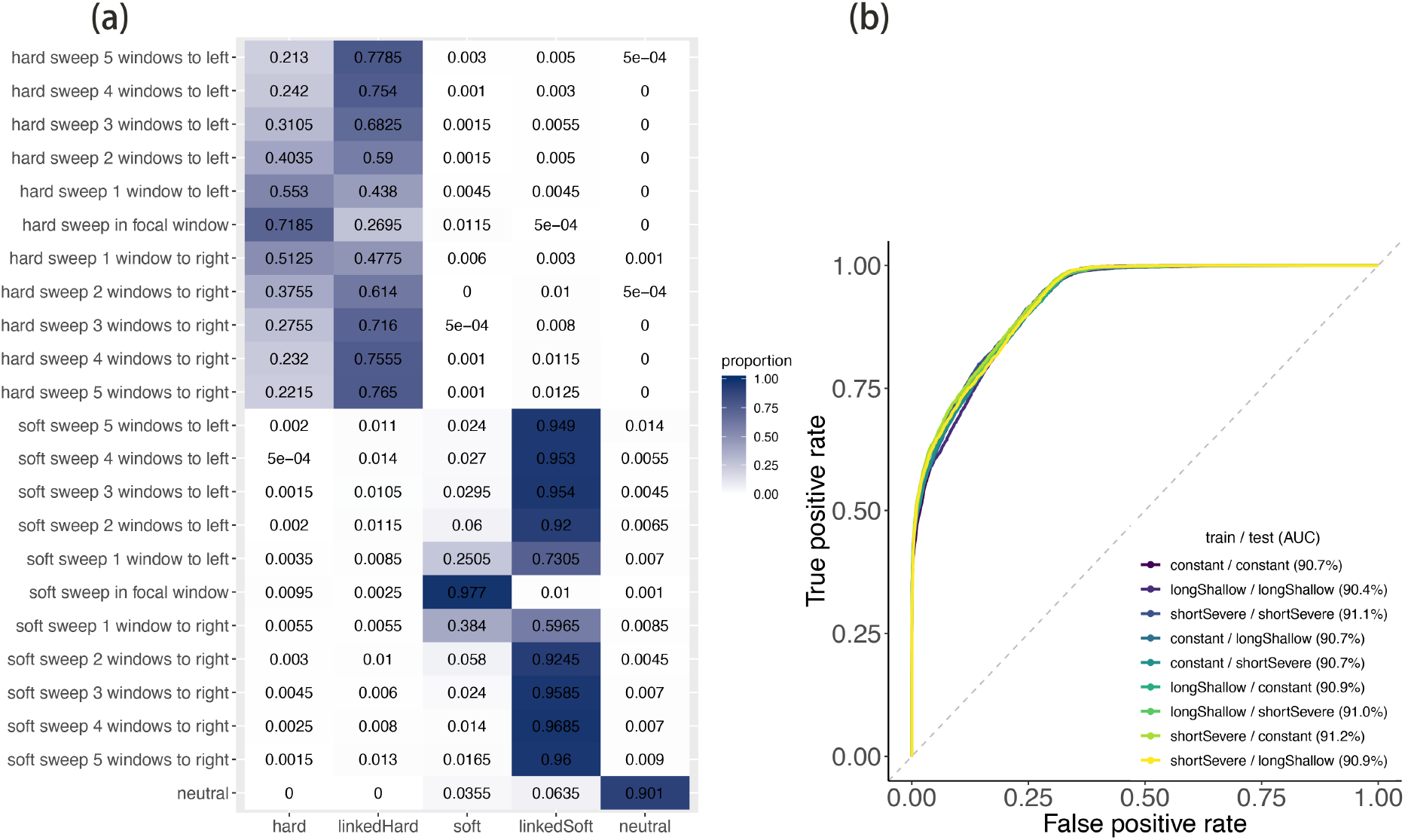
Performance of the diploS/HIC classifier on simulated data. (a) confusion matrix for a classifier trained on constant population size and validated on constant population size. (b) Shown are receiver-operator curves (ROC) for the binary classification task of hard and soft vs linked and neutral regions.

### Soft sweeps and soft-linked regions dominate the D. serrata genome

A total of 22,374 genomic regions were classified by diploS/HIC using the 5 kbp sub-window size. The results showed that linkedSoft was the most frequent classification at 46.8% (10,479 sub-windows) (**Figure 2.a**). This was followed by the neutral class, which accounted for 38.0% of all regions assessed (8,493 sub-windows) and soft sweep regions at 15.0% (3,360 sub-windows). The proportion of hard sweeps was very low at 0.1% (18 sub-windows) as was the proportion of linkedHard. We merged consecutive windows of the same sweep class and reassessed the comparison between hard and soft sweeps. There was a larger reduction in the number of distinct soft sweeps. The number of soft sweeps was reduced from 3,360 sub-windows to 2,108 clusters, whereas the number of hard sweeps was reduced from 18 sub-windows to 17 clusters. The largest sweep cluster for a soft sweep comprised 8 consecutive sub-windows which was 40kbp. Taking together all the diploS/HIC results, we observed that a large proportion of the scanned genome (62%) was impacted by soft sweeps directly or indirectly in this population.

**Figure 2.**
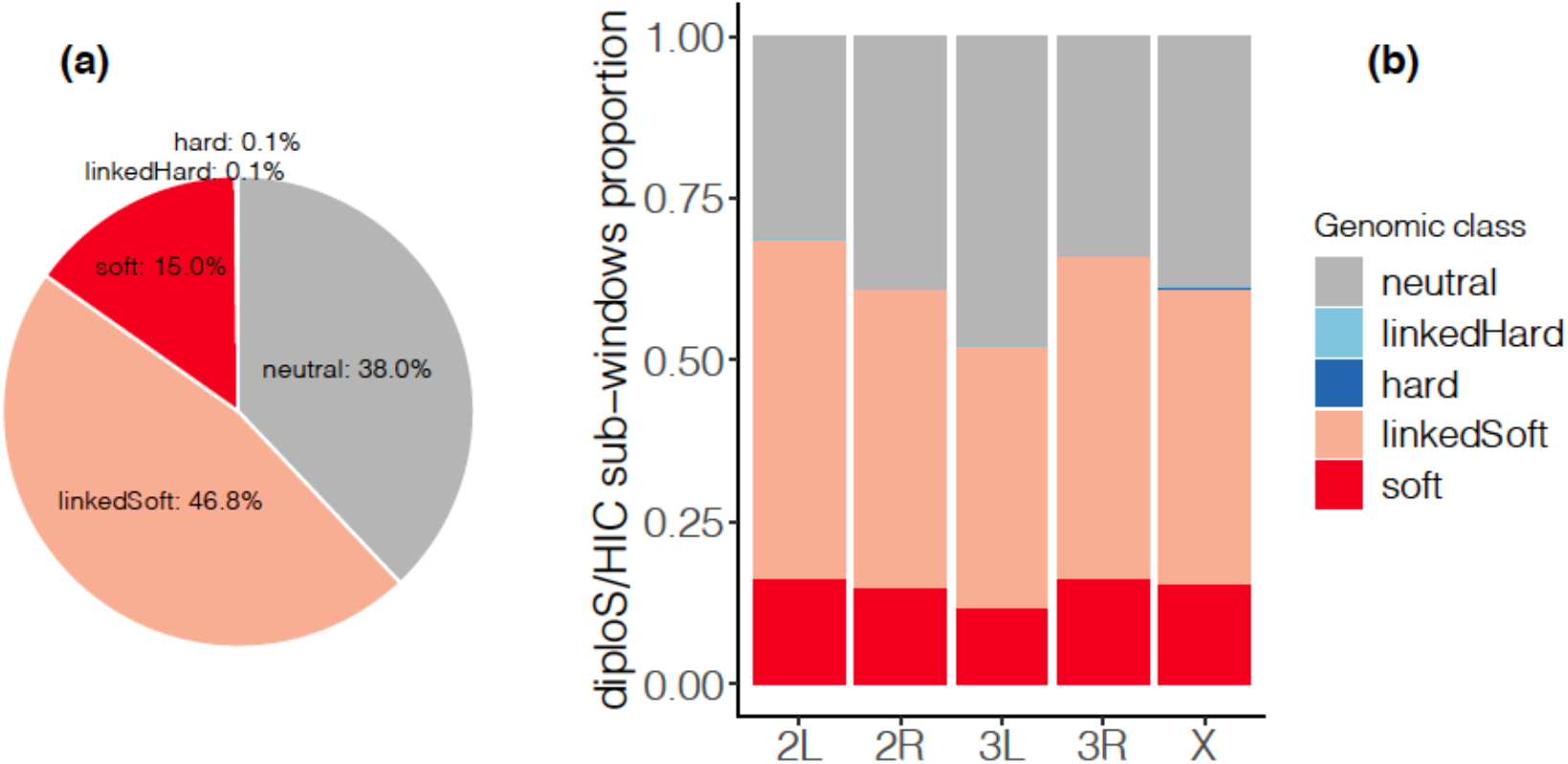
Results of diploS/HIC. (a) The percentages of five sweep classes of *D. serrata* determined by diploS/HIC. (b) The distribution of windows for each of the five sweep classes across chromosomes.

Comparison of the distribution of sweep sub-windows on different chromosome arms revealed a significant difference for soft sweeps (χ^2^=48.096, *df=4, p*=9.01×10^-10^) (**Figure 2.b**). The proportions of soft sweep sub-windows were 2L:16.36%, 2R:14.82%, 3L:11.82%, 3R:16.35% and X:15.44%. This was driven by a deficit of soft sweeps on chromosome arm 3L *(P_fdr-ajust_<0.05)*. For the 18 hard sweep sub-windows, 7 were on the X chromosome, followed by 2L (5), 3R (4), and 2L (2).

For comparison to diploS/HIC, we also searched for sweep signatures using H12 and *p*iHS. Both of the alternative methods classified far fewer selective sweeps than diploS/HIC. H12 revealed 102 selective sweeps, with largest H12 = 0.103. Most of the sweeps were likely to be soft with high H2/H1 (median H2/H1 = 0.66). *p*iHS reported 786 SNPs with *p*iHS value >= 4.08, which accounted for about 0.17% of the scanned SNPs. However, different methods have different power in different situations. For example, *p*iHS has moderate power to detect a selective sweep of intermediate frequency (30%-80%), while H12 has limited power to detect sweeps near fixation (>80%) or low frequency (<30%) (Pickrell et al., 2009; Wollstein & Stephan, 2015). Further comparison of the above three methods can be found in **Supplementary Figure S2.** We utilise the diploS/HIC results for further analysis and interpretation.

### Regions linked to soft sweeps are enriched for deleterious variants

We used snpEff and snpSift to annotate the variants within each sweep region based on their impact on the translated sequence: high, moderate, low and modifier. Of the 11,820,435 SNPs we examined, 90.34% of them were classified as ‘modifiers’; 5.93% were ‘low impact’ variants; 3.47% were ‘moderate impact’ variants and 0.27% were ‘high impact’ variants. As expected, the number of low impact variants was larger than the number of higher impact variants, even if the sweep class was taken into consideration (4 impact class x 5 sweep class contingency table: χ^2^=145560, *df*=12, *P*<2.2×10^-16^) (**Supplementary Figure S3**).

We observed a distinct distribution pattern for the impact of SNPs in different sweep classes (**Figure 3**). Based on 10,000 permutations, soft sweep regions were significantly enriched for deleterious SNPs (high, moderate, and low) by 1.61-fold, 1.51-fold and 1.42-fold, respectively. Soft sweep regions also had a deficit of modifier SNPs (0.77-fold). For linkedSoft regions, the high and moderate deleterious SNPs were significantly enriched as well (1.29-fold and 1.12-fold, respectively), with a slight deficit of modifier SNPs (0.98-fold). Although a similar pattern to soft sweeps was observed for hard sweeps, their very rare occurrence produced large confidence intervals, and thus lower statistical power.

**Figure 3.**
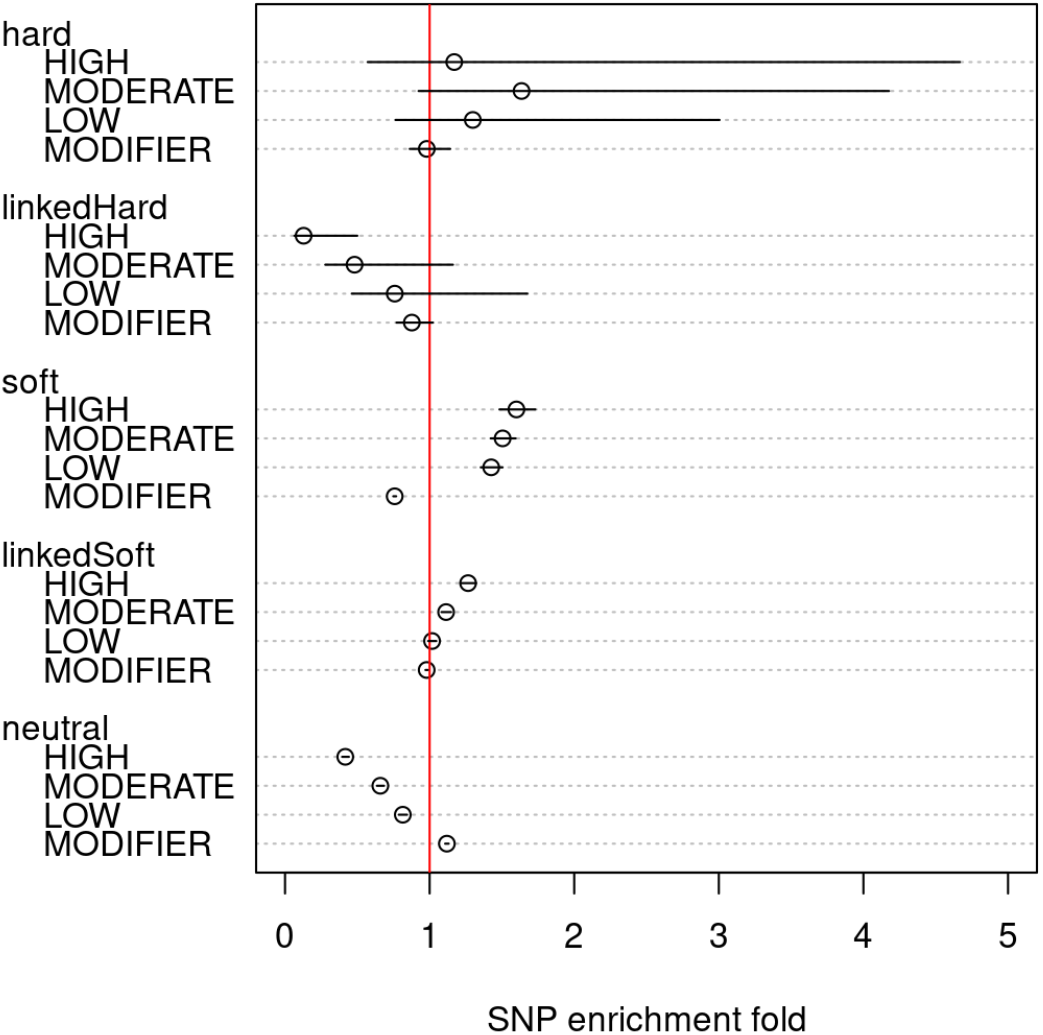
The SNP enrichment fold (open circles) for SNP effect in five sweep classes and their 95% confidence interval (black lines) compared with the permutation-based expectation (red line).

Under the assumption that variants classified by snpEff and snpSift as ‘high impact’ are more likely to have deleterious fitness effects than other classes, the enrichment of these variants in soft sweep linked regions suggests that hitchhiking may drive these alleles to higher frequencies than expected under mutation-selection-drift equilibrium. This effect was most pronounced for high impact SNPs, which were more likely to be overrepresented in linkedSoft regions (**Figure 4**). The moderate impact SNPs were also revealed to be significantly enriched on all except chromosome arm 2L.

**Figure 4.**
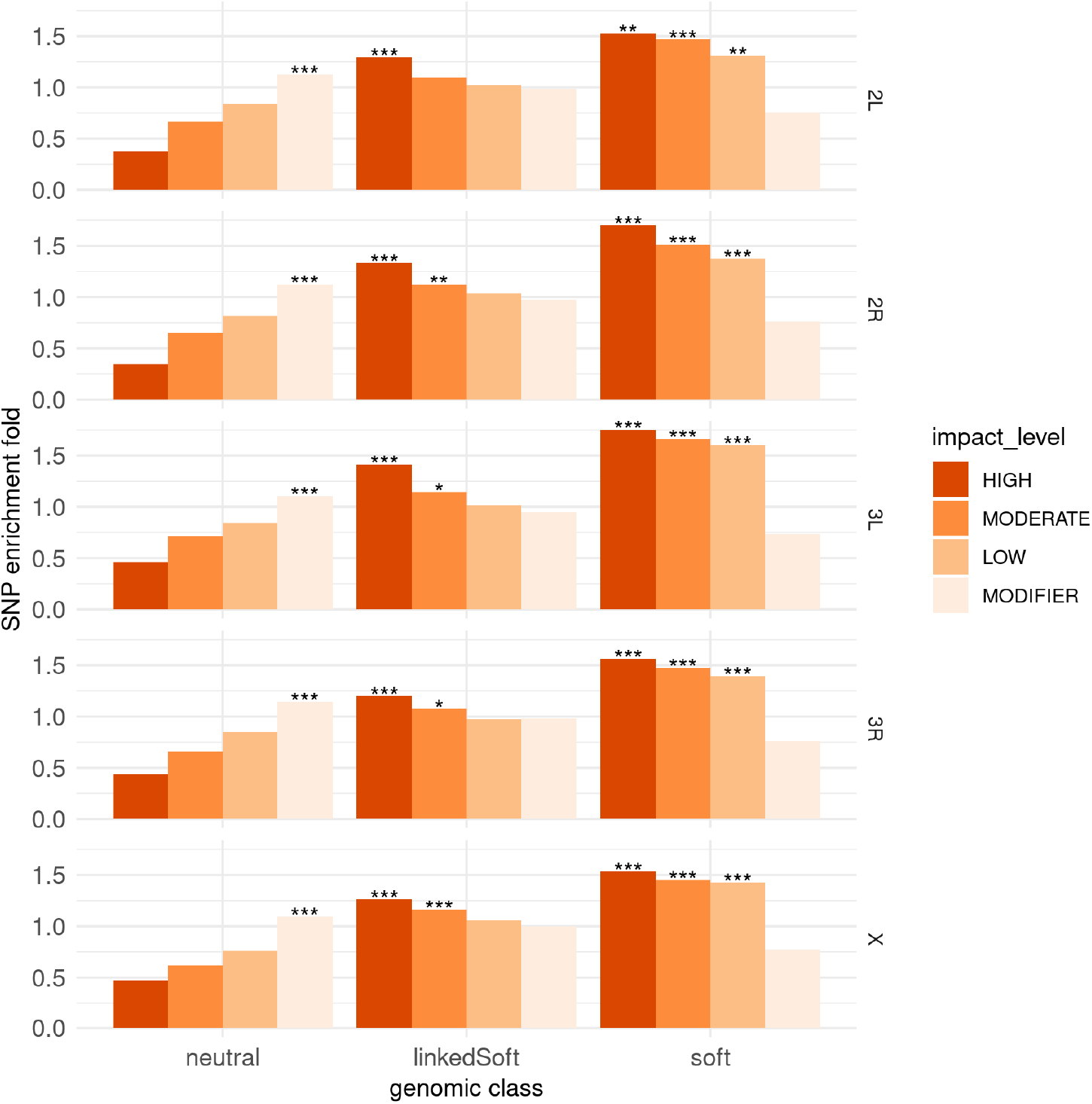
The SNP enrichment fold for SNP effect on different chromosomes. “*” indicates Bonferroni-adjusted *p* < 0.05 compared with permutation; “**” indicates Bonferroni-adjusted *p* < 0.01.

### Gene Ontology enrichment analysis

All genes within sub-windows classified as experiencing either a hard or a soft sweep were considered for gene ontology (GO) term enrichment analysis. We found 15 genes within hard sweep regions and 3,280 genes in soft sweep regions and tested the enrichment of GO terms related to these candidate genes using a permutation approach. We did not identify significantly enriched GO terms in hard sweeps with FDR-*P* < 0.05. However, in soft sweeps, a total of 323 GO terms were found to be enriched. These were comprised of 187 biological process terms, 66 molecular function terms and 70 cellular component terms, fold enrichment ranged from 1.30 to 4.08 (**Supplementary Table S1**).

Notably, some terms were involved in immune responses, such as “defence response” (GO:0006952, enrichment fold: 1.76), “defence response to fungus” (GO:0050832, 1.93). We also found some enriched GO terms related to male reproduction, including “sperm individualisation” (GO: GO:0007291, 2.03), “sperm axoneme assembly” (GO:0007288, 3.09), “male meiosis cytokinesis” (GO: GO:0007112, 2.82), “meiosis II cytokinesis” (GO:0007111, 2.76), “meiosis I cytokinesis” (GO:0007110, 3.00). Some other GO terms were enriched in the nervous system, such as “neuron projection morphogenesis” (GO:0048812, 1.62), “central nervous system development” (GO:0007417, 1.48), “neurotransmitter secretion” (GO:0007269, 1.73), “long-term memory” (GO:0007616, 1.75), “short-term memory” (GO:0007614, 2.12), “olfactory learning “(GO:0008355, 1.68), and “neurogenesis”(GO:0022008, 2.00). In addition, “oxidation-reduction process” (GO:0055114, enrichment fold: 1.53), “regulation of circadian rhythm” (GO: 0042752, 2.18), “pheromone metabolic process” (GO:0042810, 3.67), “response to caffeine” (GO:0031000, 3.52) were also recovered enriched (**Supplementary Table S1**).

### Immune-related genes and accessory gland protein genes under soft sweeps

In the soft sweep regions, we found many genes involved in immunity. *Toll* receptors are essential for immunity in *Drosophila*, playing a pivotal role in response to fungal attack, Gram-positive bacteria and virulence factors (Valanne et al., 2011). There have been nine *Toll* receptors reported in *Drosophila* (Bilak et al., 2003), and three of them were recovered in soft sweeps in this study: *Toll-6* (FBgn0036494), *Toll-7* (FBgn0034476) and *Tl* (FBgn0262473). We also found genes that were members of the *Toll* signalling pathway, including *Myd88, tube, Pellino, Spatzle-Processing Enzyme*, as well as a gene in IMD pathway: *peptidoglycan recognition protein (PGRP-LE)*.

Another set of interesting genes found in soft sweeps are those related to accessory gland protein (*Acp*) genes. Accessory gland proteins are produced by male *Drosophila* and transferred along with sperm to the female reproductive tract (Wolfner, 1997). These proteins could increase the female’s egg-laying rate, reduce female remating chance, promote sperm storage in female, be involved in sperm competition and reduce female lifespan as well (Swanson & Vacquier, 2002). In this study, *Acp62F* was recovered from soft sweeps. The soft sweep in this locus generated 40-70kb linkedSoft region on each side and a pronounced peak on H12 (>0.06) and *p*iHS (>10) (**Figure 5**).

**Figure 5.**
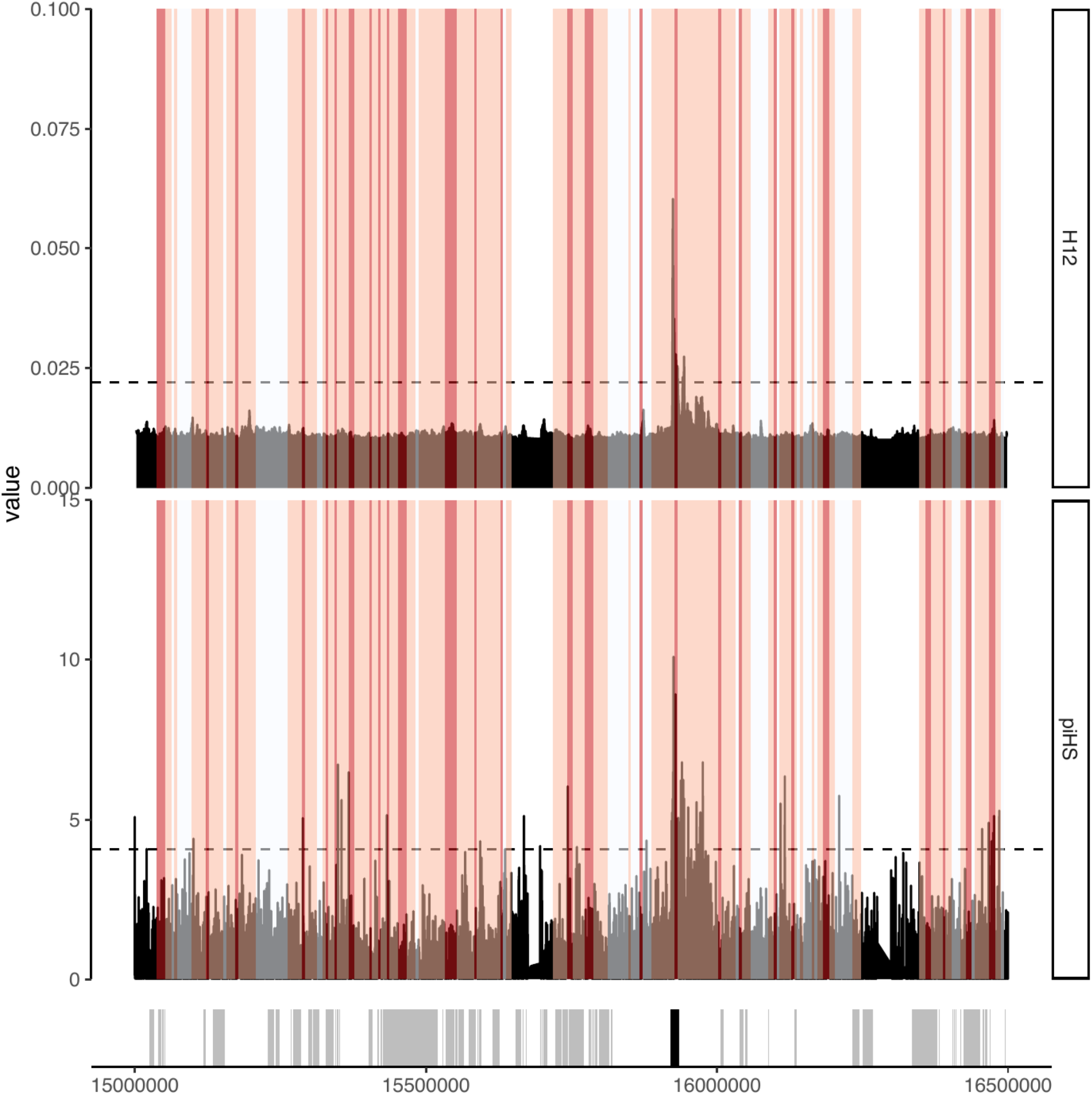
Sweep signals of *Acp62F* detected using diploS/HIC (red shaded areas), H12 and *p*iHS.Dark red – soft sweeps, light red – linkedSoft regions. Horizontal dashed lines indicate the threshold for H12 and *p*iHS. Bars on bottom panel denotes the position of genes on the chromosome (black bar: *Acp62F*, grey bars: other genes in the region).

### Prediction under bottleneck scenarios

We simulated three demographic scenarios to check the robustness of our inference – constant population size, a long shallow bottleneck followed by slight population expansion, and a short severe bottleneck. As mentioned earlier, the accuracy of prediction for diploS/HIC was very similar to the non-bottleneck scenario detailed above **(Supplementary File S1**). Although bottleneck scenarios did change the relative number of different sweep types their rank order did not change (**Figure 6**). For the long shallow bottleneck scenario followed by slight expansion, the number of sub-windows had a 10.0%-20.4% reduction depending on chromosome arm; for the short severe bottleneck scenario, the reduction ranged from between 20.7%-40.0% (**Supplementary Table S2**). In contrast, the sub-windows numbers of linkedSoft increased under the long shallow bottleneck scenario by 22.5%-36.0% and by 39.1%-54.0% under the short severe scenario. The overall genome proportion impacted directly/indirectly by soft sweeps was slightly increased from 61.9% to 73.5% (long shallow bottleneck) and 78.6% (short severe bottleneck).

**Figure 6.**
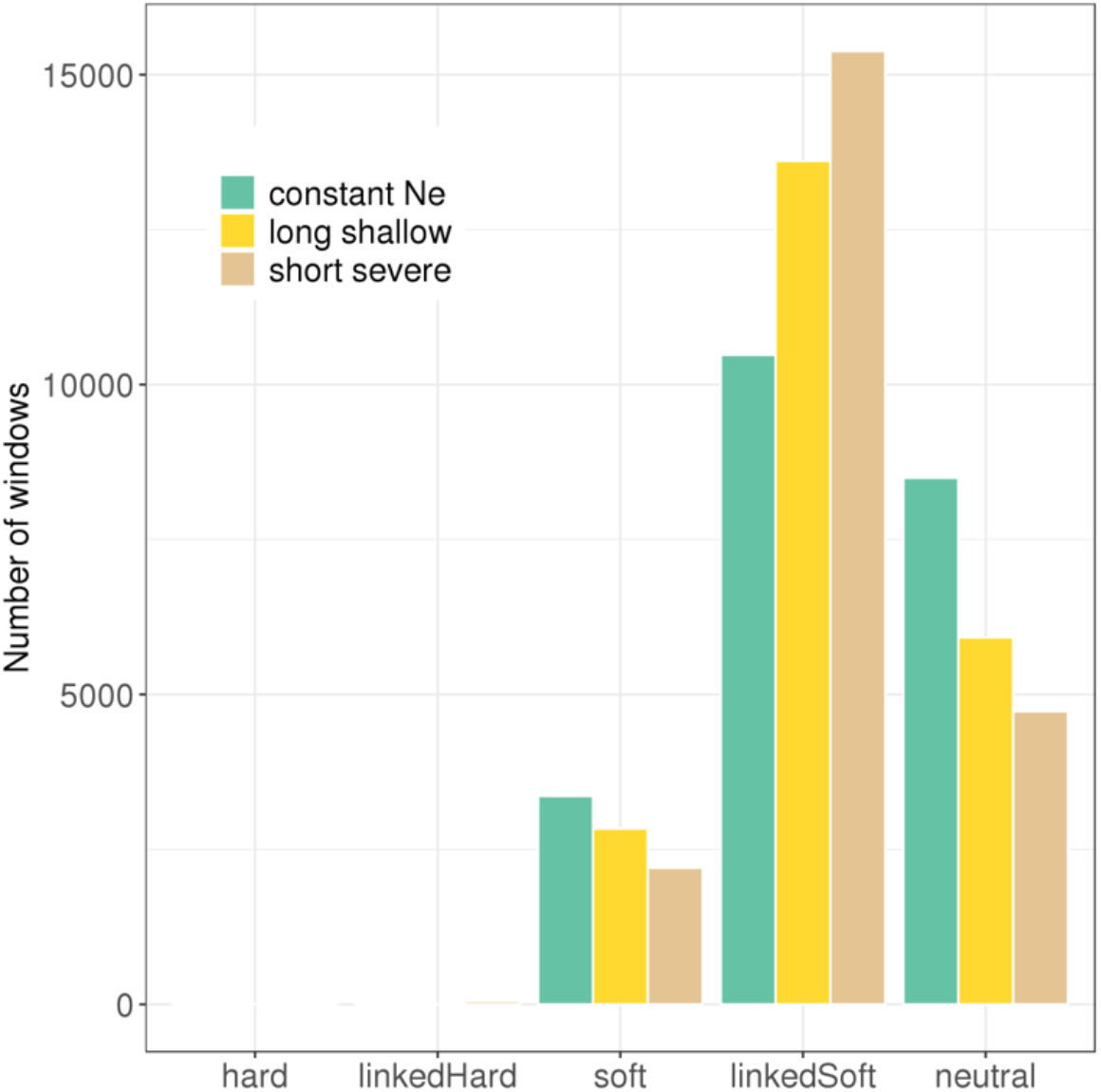
The number of genomic windows predicted by diploS/HIC under three different demographic scenarios.

We also evaluated the genes within sweep regions identified under the bottleneck scenarios. The number of candidate genes under bottleneck scenarios was reduced from 3,280 to 2,871 (long shallow bottleneck) and 2,288 (short severe bottleneck) (**Supplementary Table S2**). In addition to the three *Toll*-receptor genes *(Toll-6, Toll-7* and *Tl*), we identified another *Toll*-receptor gene (*Tehao*) in the candidate gene lists under the two bottleneck scenarios. Of the four genes involved in *Toll* signalling pathway, only *tube* was removed from the candidate gene lists. *Acp62F* remained in soft sweeps in either of the bottleneck scenarios. GO enrichment tests reported some overrepresented terms related to immunity, nervous system, and male reproduction system under the two bottleneck scenarios. Overall, the bottleneck simulations revealed that our overall conclusions were quite robust to these different demographic histories.

## Discussion

Our analysis of 110 *D. serrata* genomes provides insight into the direct and indirect impacts of positive selection on genomic variation in an endemic Drosophila population. The direct impacts, viewed through a lens of selective sweeps, indicate an abundance of soft rather than hard sweeps; a result consistent with recent adaptation being relatively unconstrained by the availability of beneficial mutations. Among the indirect impacts, we find that around half of the scanned genome is linked to selective sweeps and enriched for deleterious molecular variation. We discuss these findings and their caveats below.

### Soft sweeps dominate local adaptation in D. serrata

A key result from both our diploS/HIC and H2/H1 analyses was that soft sweeps far outnumber hard sweeps. As detection methods have improved, observation of pervasive soft sweeps is becoming increasingly common. For example, soft sweeps are abundant in *Drosophila melanogaster* (Garud et al., 2015; Garud & Petrov, 2016), *Anopheles* mosquitoes (Xue et al., 2021), several primates (Brand et al., 2021), humans (Novembre & Han, 2012; Schrider & Kern, 2017) and grasses (Bourgeois & Boissinot, 2019). A key determinant of whether hard or soft sweeps are expected is the availability of beneficial mutations at the onset of selection. Hard sweeps are expected when no beneficial variants are available for selection and the waiting time for them to arise via mutation is long (Messer & Petrov, 2013), whereas soft sweeps are expected to arise from standing variation if favourable alleles are present at the onset of selection (single-origin soft sweeps) (Pennings & Hermisson, 2006). Recurrent mutation, rather than standing variation, can also produce a soft sweep if waiting times for a beneficial mutation are very short (multiple-origin soft sweeps), due to large short-term population size and/or a high beneficial allelic mutation rate (Pennings & Hermisson, 2006). It is important to point out that our analyses are not capable of distinguishing between single- and multiple-origin soft sweeps and therefore our observation of abundant soft sweeps does not automatically imply that all recent adaptation in *D. serrata* has proceeded from standing variation (Hermisson & Pennings, 2017).

Indeed, the relevant estimates of the effective population-size (*N_e_*) scaled mutation rate (θ = *2N_e_u)* for multiple Drosophila species indicate that a combination of single and multi-origin soft sweep classes is likely (Hermisson & Pennings, 2017). For example, Watterson’s θ (Watterson, 1975) values from the most likely neutral regions of the genome, short introns, in *D. melanogaster* and *D. simulans* are 0.008 and 0.013, respectively (Andolfatto et al., 2011; Parsch et al., 2010), and if the short-term *N_e_* is underestimated by the polymorphism data by a factor of 10, then true θ is predicted to be ~0.1 (Hermisson & Pennings, 2017). With a θ value of 0.1, under most scenarios, a combination of hard sweeps and multiple origin soft sweeps is expected according to simulations by Hermisson and Pennings (2017).

We estimated θ for short introns in *D. serrata* to be ~0.2 after applying a similar adjustment of short-term *N_e_* underestimation (θ_short intron_ = 0.0217 pre-adjustment). In agreement with our empirical observation of common soft sweeps in *D. serrata*, it is expected that hard sweeps will be rare with such a θ value and that adaptation will proceed via a combination of multiple- and single-origin soft sweeps (Hermisson & Pennings, 2017). While multiple-origin soft sweeps can arise from either mutation or standing genetic variation, our θ estimate places *D. serrata* in a zone where adaptation from standing genetic variation and single-origin soft sweeps are found and potentially common (Hermisson & Pennings, 2017).

The population genetic arguments above lead us to consider how beneficial variants underlying sweeps may have existed as standing variation in the study population. One possibility is that mutations underlying soft sweeps could be maintained by fluctuating or balancing selection (Barton & Keightley, 2002). In *D. melanogaster*, traits involved host-pathogen interaction (Croze et al., 2016), mate competition (Dugand et al., 2018) and those subject to seasonal selection (Machado et al., 2021) are subject to balancing or fluctuating selection, both of which maintain variation.

Interestingly, we detected genes underlying such functions in our soft sweep windows. We identified three *Toll* receptors and many *Toll* signalling pathway factors under soft sweeps, which are involved in protection against pathogenic attack (Ferrandon et al., 2007; Valanne et al., 2011). We also observed genes related to reproduction and mate competition. For example, the accessory gland protein *Acp62F*, was indicated by both diploS/HIC and H12 and *p*iHS. As shown in **Figure 5**, there are no other genes in this genomic region making *Acp62F* a likely target of selection. *Acp62F*, protects sperm in the female reproductive tract from protease attack (Lung et al., 2002) and contributes to the up-regulation of genes involved in egg production and muscle development (Avila et al., 2011). We also note that *Acp62F* has been identified as a potential candidate for balancing selection in *D. simulans* (Begun et al., 2000).

A potential limitation of our diploS/HIC analyses concerns the unknown demographic history of *D. serrata* (Harris et al., 2018). Demographic events can leave genomic signatures similar to natural selection, such as population expansion/shrinkage or bottleneck (Akey et al., 2002; Sabeti et al., 2006; Zivkovic & Wiehe, 2008). Pervasive genome-wide negative Tajima’s D values in the study population indicate that a bottlenecked demographic history is a possibility (Reddiex et al., 2018). While training diploS/HIC on a simulated bottleneck did decrease the number of predicted sweeps relative to constant population size, the broad conclusions were unaffected. Furthermore, predictions made using mismatched demographic histories had little impact on accuracy (**Figure 1b**), suggesting that misspecifying the demographic history is not a major problem (Kern & Schrider, 2018). It is also worth noting that we have trained the classifier using a relatively recent time since fixation. As far as we know, how well diploS/HIC performs as the time since fixation increases has not been thoroughly explored, however, previous work using a similar method found that as the time since fixation increased the likelihood of misclassifying hard sweeps as soft sweeps also increased (Schrider et al., 2015) and a study that used diploS/HIC and set a very long time since fixation (0.5M years) found that diploS/HIC could not distinguish between hard and soft sweeps (Bourgeois & Boissinot, 2019). In any case, based on similar work in humans (Schrider & Kern, 2017), primates (Brand et al., 2021), and plants (Bourgeois et al., 2018), which used a much longer time since fixation and still found many more soft sweeps then hard sweeps, we do not expect the potential underestimate of the number of hard sweeps to be considerable.

### Indirect impacts of positive selection on genomic variation

One of the theoretical consequences resulting from selective sweeps is the elevated frequency of deleterious variation via hitchhiking (Smith & Haigh, 1974). In these regions, purifying selection cannot work efficiently to remove deleterious variants from the population, particularly in regions of low recombination (Hadany & Feldman, 2005; Hartfield & Otto, 2011). In this study, soft and soft sweep-linked regions were widespread and significantly enriched for detrimental SNPs, suggesting that hitchhiking may have indeed maintained deleterious variation above what is expected under mutation-selection balance. While hard sweeps also had a trend towards enrichment for detrimental SNPs, this result was far less conclusive due to the low number of hard sweeps and low statistical power. Perhaps not unexpectedly, there was weaker but still significant, overrepresentation of deleterious variation in soft-linked sweeps than within soft sweep region themselves (~1.15 fold compared to ~1.50 fold). Similar effects have been observed in human studies (Schrider & Kern, 2017). Although the enrichment fold could be considered marginal, the overall impact of linked deleterious variation may not be negligible given such a large fraction of the genome may be linked to soft sweeps. An interesting consideration is that linked selection may in fact increase the likelihood of future adaptation via standing variation because some deleterious variants may become beneficial in the face of a shift in selective optima.

### Conclusion

Machine learning approaches such as diploS/HIC are becoming increasingly popular for population genomic analysis. The ability to scan and classify the whole genome provides intuitive insights into the possible direct and indirect impacts of selection on a genome-wide scale. While powerful, it is likely that further refinements are needed. For example, partial sweeps and balancing selection could be incorporated into new classifiers. Our study suggests that it may be fruitful to consider the interplay between fluctuating selection and sweeps. Finally, it is important to note that while our study considers fixation events at individual loci, GWAS underscore the reality that variation in most phenotypic traits is controlled by many genes, which raises the possibility that selection is often polygenic (Barghi et al., 2020; Pritchard & Di Rienzo, 2010; Pritchard et al., 2010). This would still point to a significant role for standing as opposed to mutational variation in response to positive selection, it may be the case that what we can detect here is an underestimate of the selection occurring on standing variation in nature.

## Acknowledgements

We thank T. Flatt, M. Kapun, D. Ortiz Barrientos for comments on early drafts of this work. This research was supported by funding from the Australian Research Council and the University of Queensland.

## Data Accessibility

All genome sequence data are available at NCBI Bioproject ID: PRJNA419238

## Author Contributions

S.C., Y.W. and S. A. designed the study; Y.W. performed the analyses. A.R. conducted the fly experiments, sequencing, alignments and SNP calling; Y.W., S.A. and S.C. wrote the manuscript.

## References

Akey, J. M., Zhang, G., Zhang, K., Jin, L., & Shriver, M. D. (2002). Interrogating a high-density SNP map for signatures of natural selection. Genome Research, 12(12), 1805–1814. doi:10.1101/gr.631202

Allen, S. L., Bonduriansky, R., Sgro, C. M., & Chenoweth, S. F. (2017). Sex-biased transcriptome divergence along a latitudinal gradient. Molecular Ecology, 26(5), 1256–1272. doi:10.1111/mec.14015

Allen, S. L., Delaney, E. K., Kopp, A., & Chenoweth, S. F. (2017). Single-molecule sequencing of the *Drosophila serrata* genome. G3: Genes, Genomes, Genetics, 7(3), 781–788. doi:10.1534/g3.116.037598

Andolfatto, P., Wong, K. M., & Bachtrog, D. (2011). Effective Population Size and the Efficacy of Selection on the X Chromosomes of Two Closely Related Drosophila Species. Genome Biology and Evolution, 3, 114–128. doi:10.1093/gbe/evq086

Avila, F. W., Sirot, L. K., LaFlamme, B. A., Rubinstein, C. D., & Wolfner, M. F. (2011). Insect seminal fluid proteins: identification and function. Annual Review of Entomology, 56, 21–40. doi:10.1146/annurev-ento-120709-144823

Barghi, N., Hermisson, J., & Schlotterer, C. (2020). Polygenic adaptation: a unifying framework to understand positive selection (vol 18, pg 913, 2020). Nature Reviews Genetics, 21(12), 782–782. doi:10.1038/s41576-020-0276-2

Barrett, R. D., & Schluter, D. (2008). Adaptation from standing genetic variation. Trends in Ecology & Evolution, 23(1), 38–44. doi:10.1016/j.tree.2007.09.008

Barton, N. H. (1998). The effect of hitch-hiking on neutral genealogies. Genetics Research, 72(2), 123–133. doi:Doi 10.1017/S0016672398003462

Barton, N. H., & Keightley, P. D. (2002). Understanding quantitative genetic variation. Nature Reviews Genetics, 3(1), 11–21.

Begun, D. J., Whitley, P., Todd, B. L., Waldrip-Dail, H. M., & Clark, A. G. (2000). Molecular population genetics of male accessory gland proteins in drosophila. Genetics, 156(4), 1879–1888.

Bilak, H., Tauszig-Delamasure, S., & Imler, J. L. (2003). Toll and Toll-like receptors in *Drosophila*. Biochemical Society Transactions, 31(Pt 3), 648–651. doi:10.1042/bst0310648

Blows, M. W., & Hoffmann, A. A. (1993). The genetics of central and marginal populations of Drosophila serrata. I. Genetic variation for stress resistance and species borders. Evolution, 47(4), 1255–1270.

Bourgeois, Y., & Boissinot, S. (2019). Selection at behavioural, developmental and metabolic genes is associated with the northward expansion of a successful tropical colonizer. Molecular Ecology, 28(15), 3523–3543. doi:10.1111/mec.15162

Bourgeois, Y., Stritt, C., Walser, J. C., Gordon, S. P., Vogel, J. P., & Roulin, A. C. (2018). Genome-wide scans of selection highlight the impact of biotic and abiotic constraints in natural populations of the model grass Brachypodium distachyon. Plant Journal, 96(2), 438–451. doi:10.1111/tpj.14042

Brand, C. M., White, F. J., Ting, N., & Webster, T. H. (2021). Soft sweeps predominate recent positive selection in bonobos (Pan paniscus) and chimpanzees (Pan troglodytes). bioRxiv, 2020.2012.2014.422788. doi:10.1101/2020.12.14.422788

Chenoweth, S. F., & Blows, M. W. (2003). Signal trait sexual dimorphism and mutual sexual selection in *Drosophila serrata*. Evolution, 57(10), 2326–2334.

Cingolani, P., Platts, A., Wang le, L., Coon, M., Nguyen, T., Wang, L., … Ruden, D. M. (2012). A program for annotating and predicting the effects of single nucleotide polymorphisms, SnpEff: SNPs in the genome of *Drosophila melanogaster* strain w1118; iso-2; iso-3. Fly (Austin), 6(2), 80–92. doi:10.4161/fly.19695

Colosimo, P. F., Hosemann, K. E., Balabhadra, S., Villarreal, G., Jr., Dickson, M., Grimwood, J., … Kingsley, D. M. (2005). Widespread parallel evolution in sticklebacks by repeated fixation of Ectodysplasin alleles. Science, 307(5717), 1928–1933. doi:10.1126/science.1107239

Croucher, N. J., Chewapreecha, C., Hanage, W. P., Harris, S. R., McGee, L., van der Linden, M., … Bentley, S. D. (2014). Evidence for soft selective sweeps in the evolution of pneumococcal multidrug resistance and vaccine escape. Genome Biology Evolution, 6(7), 1589–1602. doi:10.1093/gbe/evu120

Croze, M., Živković, D., Stephan, W., & Hutter, S. (2016). Balancing selection on immunity genes: review of the current literature and new analysis in *Drosophila melanogaster*. Zoology, 119(4), 322–329.

Dugand, R. J., Kennington, W. J., & Tomkins, J. L. (2018). Evolutionary divergence in competitive mating success through female mating bias for good genes. Science advances, 4(5), eaaq0369.

Durrett, R., & Schweinsberg, J. (2004). Approximating selective sweeps. Theoretical Population Biology, 66(2), 129–138. doi:10.1016/j.tpb.2004.04.002

Falconer, D. S. (1996). Introduction to quantitative genetics: Pearson Education India.

Fay, J. C., & Wu, C.-I. (2000). Hitchhiking under positive Darwinian selection. Genetics, 155(3), 1405–1413.

Ferrandon, D., Imler, J. L., Hetru, C., & Hoffmann, J. A. (2007). The *Drosophila* systemic immune response: sensing and signalling during bacterial and fungal infections. Nature Reviews Immunology, 7(11), 862–874. doi:10.1038/nri2194

Frentiu, F. D., Adamski, M., McGraw, E. A., Blows, M. W., & Chenoweth, S. F. (2009). An expressed sequence tag (EST) library for Drosophila serrata, a model system for sexual selection and climatic adaptation studies. BMC Genomics, 10, 40. doi:10.1186/1471-2164-10-40

Garud, N. R., Messer, P. W., Buzbas, E. O., & Petrov, D. A. (2015). Recent selective sweeps in North American *Drosophila melanogaster* show signatures of soft sweeps. Plos Genetics, 11(2), e1005004.

Garud, N. R., & Petrov, D. A. (2016). Elevated linkage disequilibrium and signatures of soft sweeps are common in *Drosophila melanogaster*. Genetics, 203(2), 863–880. doi:10.1534/genetics.115.184002

Gautier, M., & Naves, M. (2011). Footprints of selection in the ancestral admixture of a New World Creole cattle breed. Molecular Ecology, 20(15), 3128–3143. doi:10.1111/j.1365-294X.2011.05163.x

Gautier, M., & Vitalis, R. (2012). rehh: an R package to detect footprints of selection in genome-wide SNP data from haplotype structure. Bioinformatics, 28(8), 1176–1177. doi:10.1093/bioinformatics/bts115

Gosden, T. P., Reddiex, A. J., & Chenoweth, S. F. (2018). Artificial selection reveals sex differences in the genetic basis of sexual attractiveness. Proceedings of the National Academy of Sciences, 115(21), 5498–5503. doi:10.1073/pnas.1720368115

Haag-Liautard, C., Dorris, M., Maside, X., Macaskill, S., Halligan, D. L., Houle, D., … Keightley, P. D. (2007). Direct estimation of per nucleotide and genomic deleterious mutation rates in *Drosophila*. Nature, 445(7123), 82–85. doi:10.1038/nature05388

Hadany, L., & Feldman, M. W. (2005). Evolutionary traction: the cost of adaptation and the evolution of sex. Journal of Evolutionary Biology, 18(2), 309–314. doi:10.1111/j.1420-9101.2004.00858.x

Harris, R. B., Sackman, A., & Jensen, J. D. (2018). On the unfounded enthusiasm for soft selective sweeps II: Examining recent evidence from humans, flies, and viruses. Plos Genetics, 14(12), e1007859. doi:10.1371/journal.pgen.1007859

Hartfield, M., & Otto, S. P. (2011). Recombination and Hitchhiking of Deleterious Alleles. Evolution, 65(9), 2421–2434. doi:10.1111/j.1558-5646.2011.01311.x

Hermisson, J., & Pennings, P. S. (2005). Soft sweeps: molecular population genetics of adaptation from standing genetic variation. Genetics, 169(4), 2335–2352. doi:10.1534/genetics.104.036947

Hermisson, J., & Pennings, P. S. (2017). Soft sweeps and beyond: understanding the patterns and probabilities of selection footprints under rapid adaptation. Methods in Ecology and Evolution, 8(6), 700–716.

Hine, E., Chenoweth, S. F., & Blows, M. W. (2004). Multivariate quantitative genetics and the lek paradox: genetic variance in male sexually selected traits of *Drosophila serrata* under field conditions. Evolution, 58(12), 2754–2762.

Hine, E., Lachish, S., Higgie, M., & Blows, M. W. (2002). Positive genetic correlation between female preference and offspring fitness. Proceedings of the Royal Society of London. Series B: Biological Sciences, 269(1506), 2215–2219.

Innan, H., & Kim, Y. (2004). Pattern of polymorphism after strong artificial selection in a domestication event. Proceedings of the National Academy of Sciences, 101(29), 10667–10672. doi:10.1073/pnas.0401720101

Jenkins, N. L., & Hoffmann, A. A. (2001). Distribution of *Drosophila serrata Malloch* (*Diptera: Drosophilidae*) in Australia with particular reference to the southern border. Australian Journal of Entomology, 40(1), 41–48.

Jensen, J. D. (2014). On the unfounded enthusiasm for soft selective sweeps. Nature Communication, 5, 5281. doi:10.1038/ncomms6281

Jones, B. L., Raga, T. O., Liebert, A., Zmarz, P., Bekele, E., Danielsen, E. T., … Swallow, D. M. (2013). Diversity of lactase persistence alleles in Ethiopia: signature of a soft selective sweep. The American Journal of Human Genetics, 93(3), 538–544. doi:10.1016/j.ajhg.2013.07.008

Kaplan, N. L., Hudson, R. R., & Langley, C. H. (1989). The hitchhiking effect revisited. Genetics, 123(4), 887–899.

Keightley, P. D., Ness, R. W., Halligan, D. L., & Haddrill, P. R. (2014). Estimation of the spontaneous mutation rate per nucleotide site in a *Drosophila melanogaster* full-sib family. Genetics, 196(1), 313–320. doi:10.1534/genetics.113.158758

Kelly, J. K. (1997). A test of neutrality based on interlocus associations. Genetics, 146(3), 1197–1206.

Kern, A. D., & Schrider, D. R. (2016). Discoal: flexible coalescent simulations with selection. Bioinformatics, 32(24), 3839–3841. doi:10.1093/bioinformatics/btw556

Kern, A. D., & Schrider, D. R. (2018). diploS/HIC: An updated approach to classifying selective sweeps. G3: Genes, Genomes, Genetics, 8(6), 1959–1970. doi:10.1534/g3.118.200262

Li, H. (2013). Aligning sequence reads, clone sequences and assembly contigs with BWA-MEM. arXiv preprint arXiv:1303.3997.

Lung, O., Tram, U., Finnerty, C. M., Eipper-Mains, M. A., Kalb, J. M., & Wolfner, M. F. (2002). The *Drosophila melanogaster* seminal fluid protein Acp62F is a protease inhibitor that is toxic upon ectopic expression. Genetics, 160(1), 211–224.

Lush, J. L. (2013). Animal breeding plans: Read Books Ltd.

Lynch, M., & Walsh, B. (1998). Genetics and analysis of quantitative traits.

Machado, H. E., Bergland, A. O., Taylor, R., Tilk, S., Behrman, E., Dyer, K., … Karasov, T. L. (2021). Broad geographic sampling reveals the shared basis and environmental correlates of seasonal adaptation in Drosophila. Elife, 10, e67577.

McKenna, A., Hanna, M., Banks, E., Sivachenko, A., Cibulskis, K., Kernytsky, A., … DePristo, M. A. (2010). The Genome Analysis Toolkit: A MapReduce framework for analyzing next-generation DNA sequencing data. Genome Research, 20(9), 1297–1303. doi:10.1101/gr.107524.110

Messer, P. W., & Petrov, D. A. (2013). Population genomics of rapid adaptation by soft selective sweeps. Trends in Ecology & Evolution, 28(11), 659–669. doi:10.1016/j.tree.2013.08.003

Nielsen, R. (2005). Molecular signatures of natural selection. Annual review of genetics, 39, 197–218. doi:10.1146/annurev.genet.39.073003.112420

Novembre, J., & Han, E. J. (2012). Human population structure and the adaptive response to pathogen-induced selection pressures. Philosophical Transactions of the Royal Society B-Biological Sciences, 367(1590), 878–886. doi:10.1098/rstb.2011.0305

Parsch, J., Novozhilov, S., Saminadin-Peter, S. S., Wong, K. M., & Andolfatto, P. (2010). On the Utility of Short Intron Sequences as a Reference for the Detection of Positive and Negative Selection in Drosophila. Molecular biology and evolution, 27(6), 1226–1234. doi:10.1093/molbev/msq046

Pennings, P. S., & Hermisson, J. (2006). Soft sweeps II—molecular population genetics of adaptation from recurrent mutation or migration. Molecular biology and evolution, 23(5), 1076–1084.

Pickrell, J. K., Coop, G., Novembre, J., Kudaravalli, S., Li, J. Z., Absher, D., … Pritchard, J. K. (2009). Signals of recent positive selection in a worldwide sample of human populations. Genome Research, 19(5), 826–837. doi:10.1101/gr.087577.108

Pritchard, J. K., & Di Rienzo, A. (2010). Adaptation - not by sweeps alone. Nature Reviews Genetics, 11(10), 665–667. doi:DOI 10.1038/nrg2880

Pritchard, J. K., Pickrell, J. K., & Coop, G. (2010). The Genetics of Human Adaptation: Hard Sweeps, Soft Sweeps, and Polygenic Adaptation. Current Biology, 20(4), R208–R215. doi:10.1016/j.cub.2009.11.055

Przeworski, M., Coop, G., & Wall, J. D. (2005). The signature of positive selection on standing genetic variation. Evolution, 59(11), 2312–2323.

Raquin, A. L., Brabant, P., Rhone, B., Balfourier, F., Leroy, P., & Goldringer, I. (2008). Soft selective sweep near a gene that increases plant height in wheat. Molecular Ecology, 17(3), 741–756. doi:10.1111/j.1365-294X.2007.03620.x

Reddiex, A. J., Allen, S. L., & Chenoweth, S. F. (2018). A Genomic Reference Panel for *Drosophila serrata*. G3: Genes, Genomes, Genetics, 8(4), 1335–1346. doi:10.1534/g3.117.300487

Ruden, D. M., Cingolani, P., Patel, V. M., Coon, M., Nguyen, T., Land, S. J., & Lu, X. (2012). Using *Drosophila melanogaster* as a model for genotoxic chemical mutational studies with a new program, SnpSift. Frontiers in Genetics, 3, 35.

Rundle, H. D., Chenoweth, S. F., Doughty, P., & Blows, M. W. (2005). Divergent selection and the evolution of signal traits and mating preferences. PLoS Biology, 3(11), 1988–1995. doi:10.1371/journal.pbio.0030368

Sabeti, P. C., Schaffner, S. F., Fry, B., Lohmueller, J., Varilly, P., Shamovsky, O., … Lander, E. S. (2006). Positive natural selection in the human lineage. Science, 312(5780), 1614–1620. doi:10.1126/science.1124309

Schrider, D. R., & Kern, A. D. (2017). Soft sweeps are the dominant mode of adaptation in the human genome. Molecular Biology and Evolution. doi:10.1093/molbev/msx154

Schrider, D. R., Mendes, F. K., Hahn, M. W., & Kern, A. D. (2015). Soft shoulders ahead: spurious signatures of soft and partial selective sweeps result from linked hard sweeps. Genetics, 200(1), 267–284. doi:10.1534/genetics.115.174912

Sinha, P., Dincer, A., Virgil, D., Xu, G., Poh, Y. P., & Jensen, J. D. (2011). On detecting selective sweeps using single genomes. Frontiers in Genetics, 2, 85. doi:10.3389/fgene.2011.00085

Smit, AFA, Hubley, & P, R. G. (1996-2010). RepeatMasker Open-3.0. http://www.repeatmasker.org.

Smith, J. M., & Haigh, J. (1974). The hitch-hiking effect of a favourable gene. Genetics Research, 23(1), 23–35.

Stone, E. A. (2012). Joint genotyping on the fly: identifying variation among a sequenced panel of inbred lines. Genome Research, 22(5), 966–974.

Swanson, W. J., & Vacquier, V. D. (2002). The rapid evolution of reproductive proteins. Nature Reviews Genetics, 3(2), 137–144. doi:10.1038/nrg733

Tajima, F. (1983). Evolutionary relationship of DNA sequences in finite populations. Genetics, 105(2), 437–460.

Tajima, F. (1989). Statistical method for testing the neutral mutation hypothesis by DNA polymorphism. Genetics, 123(3), 585–595.

Urbinati, I., Stafuzza, N. B., Oliveira, M. T., Chud, T. C., Higa, R. H., Regitano, L. C., … Munari, D. P. (2016). Selection signatures in Canchim beef cattle. Journal of Animal Science and Biotechnology, 7, 29. doi:10.1186/s40104-016-0089-5

Valanne, S., Wang, J. H., & Ramet, M. (2011). The *Drosophila* Toll signaling pathway. The Journal of Immunology, 186(2), 649–656. doi:10.4049/jimmunol.1002302

Voight, B. F., Kudaravalli, S., Wen, X., & Pritchard, J. K. (2006). A map of recent positive selection in the human genome. PLoS Biology, 4(3), e72. doi:10.1371/journal.pbio.0040072

Watterson, G. (1975). On the number of segregating sites in genetical models without recombination. Theoretical Population Biology, 7(2), 256–276.

Wilson, B. A., Pennings, P. S., & Petrov, D. A. (2017). Soft selective sweeps in evolutionary rescue. Genetics, 205(4), 1573–1586. doi:10.1534/genetics.116.191478

Wilson, B. A., Petrov, D. A., & Messer, P. W. (2014). Soft selective sweeps in complex demographic scenarios. Genetics, 198(2), 669–684. doi:10.1534/genetics.114.165571

Wolfner, M. F. (1997). Tokens of love: Functions and regulation of *Drosophila* male accessory gland products. Insect Biochemistry and Molecular Biology, 27(3), 179–192. doi:Doi 10.1016/S0965-1748(96)00084-7

Wollstein, A., & Stephan, W. (2015). Inferring positive selection in humans from genomic data. Investigative Genetics, 6, 5. doi:10.1186/s13323-015-0023-1

Xue, A. T., Schrider, D. R., & Kern, A. D. (2021). Discovery of ongoing selective sweeps within Anopheles mosquito populations using deep learning. Molecular biology and evolution, 38(3), 1168–1183.

Zivkovic, D., & Wiehe, T. (2008). Second-order moments of segregating sites under variable population size. Genetics, 180(1), 341–357. doi:10.1534/genetics.108.091231

