## Supplementary figures and images for "Direct and indirect impacts of positive selection on genomic variation in *Drosophila serrata*"

### Supplemental Figure 1

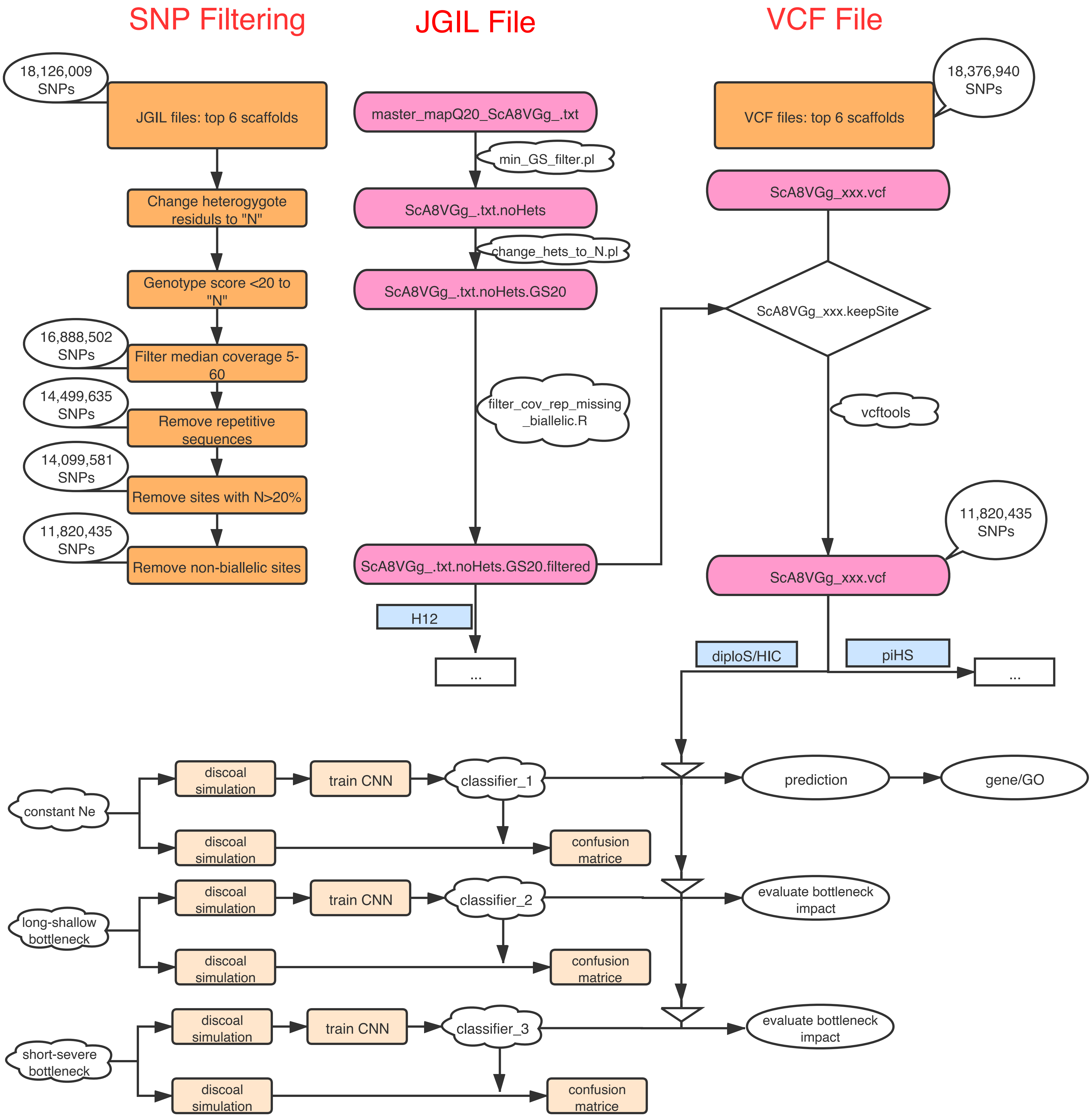

### Supplemental Figure 2

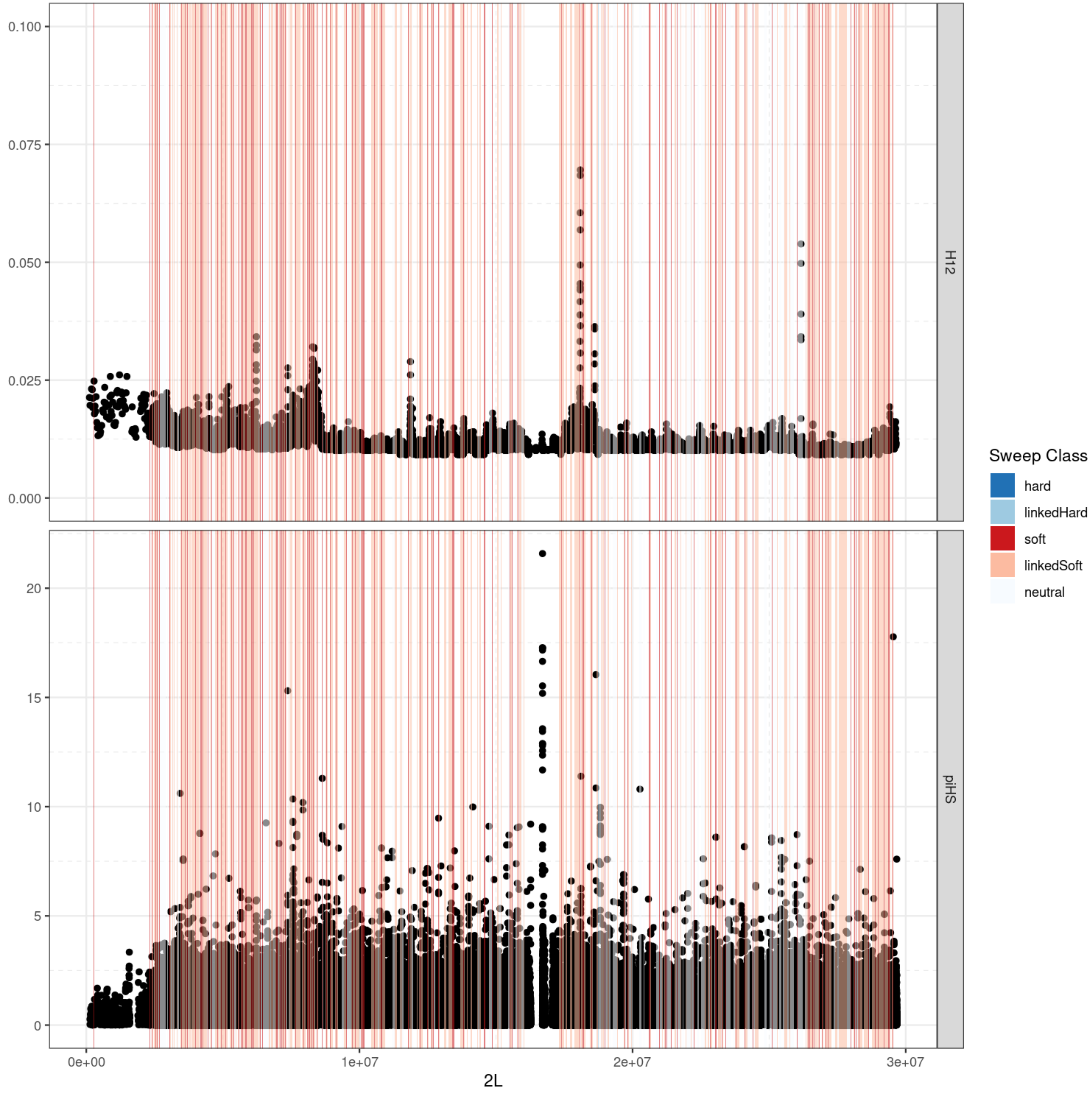

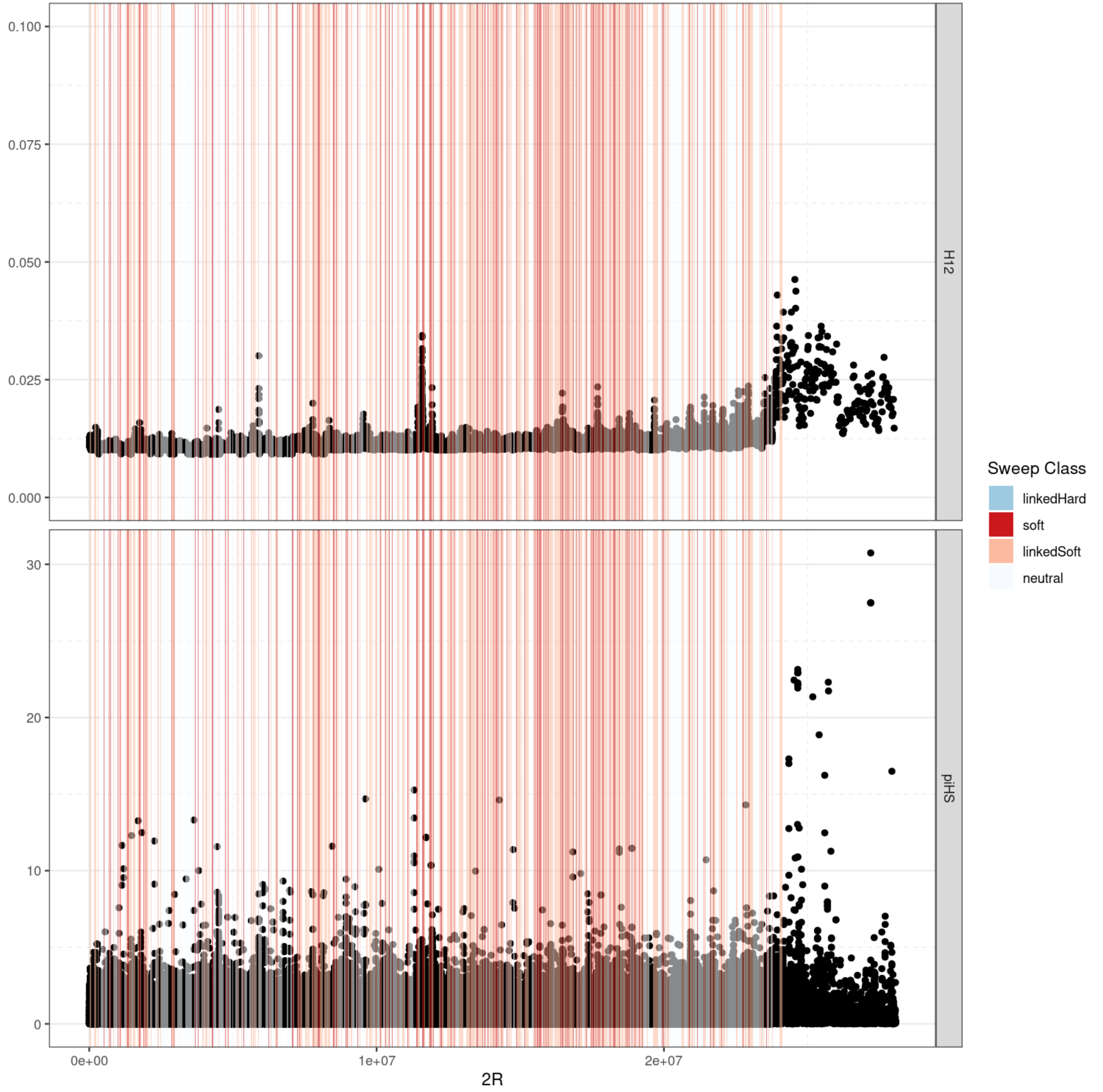

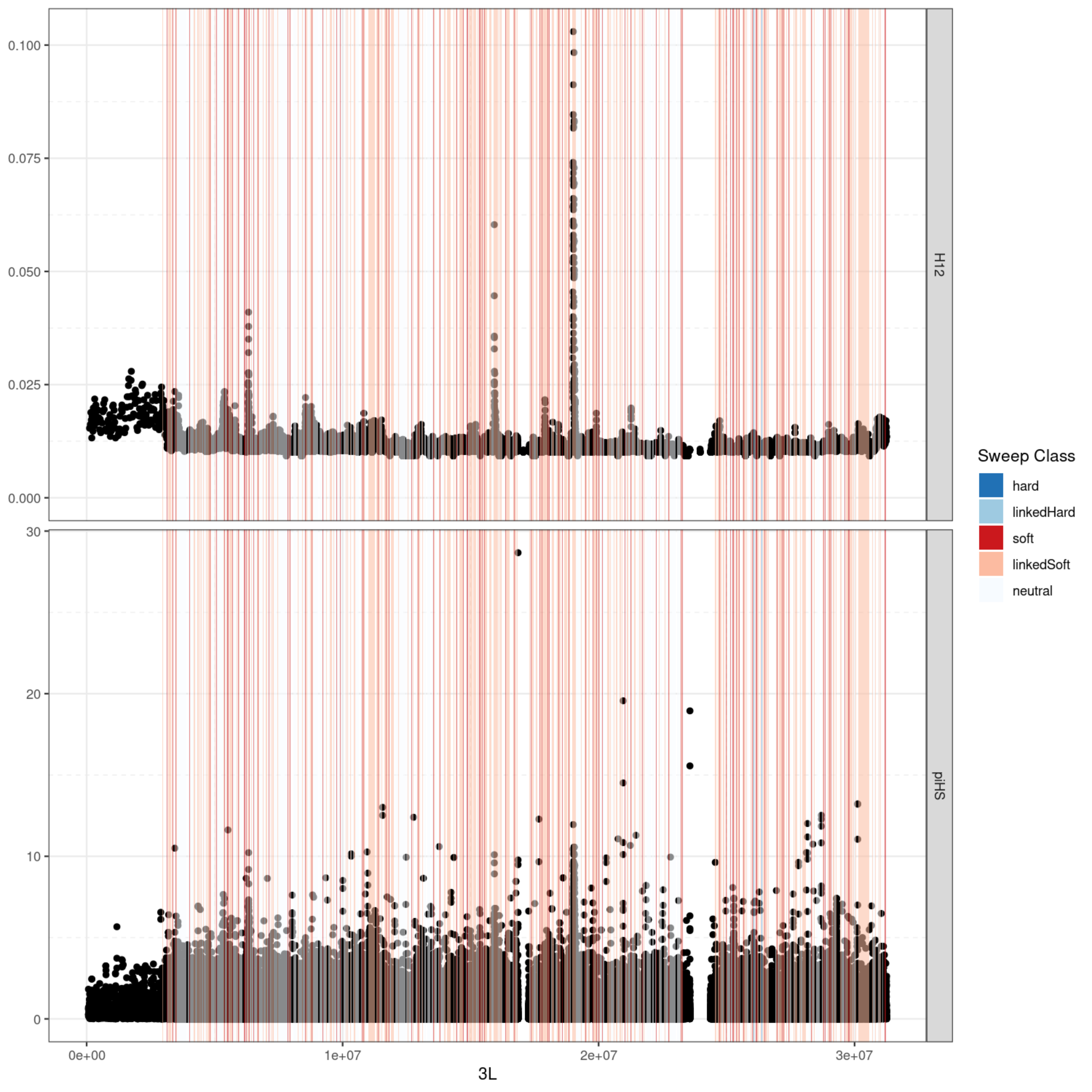

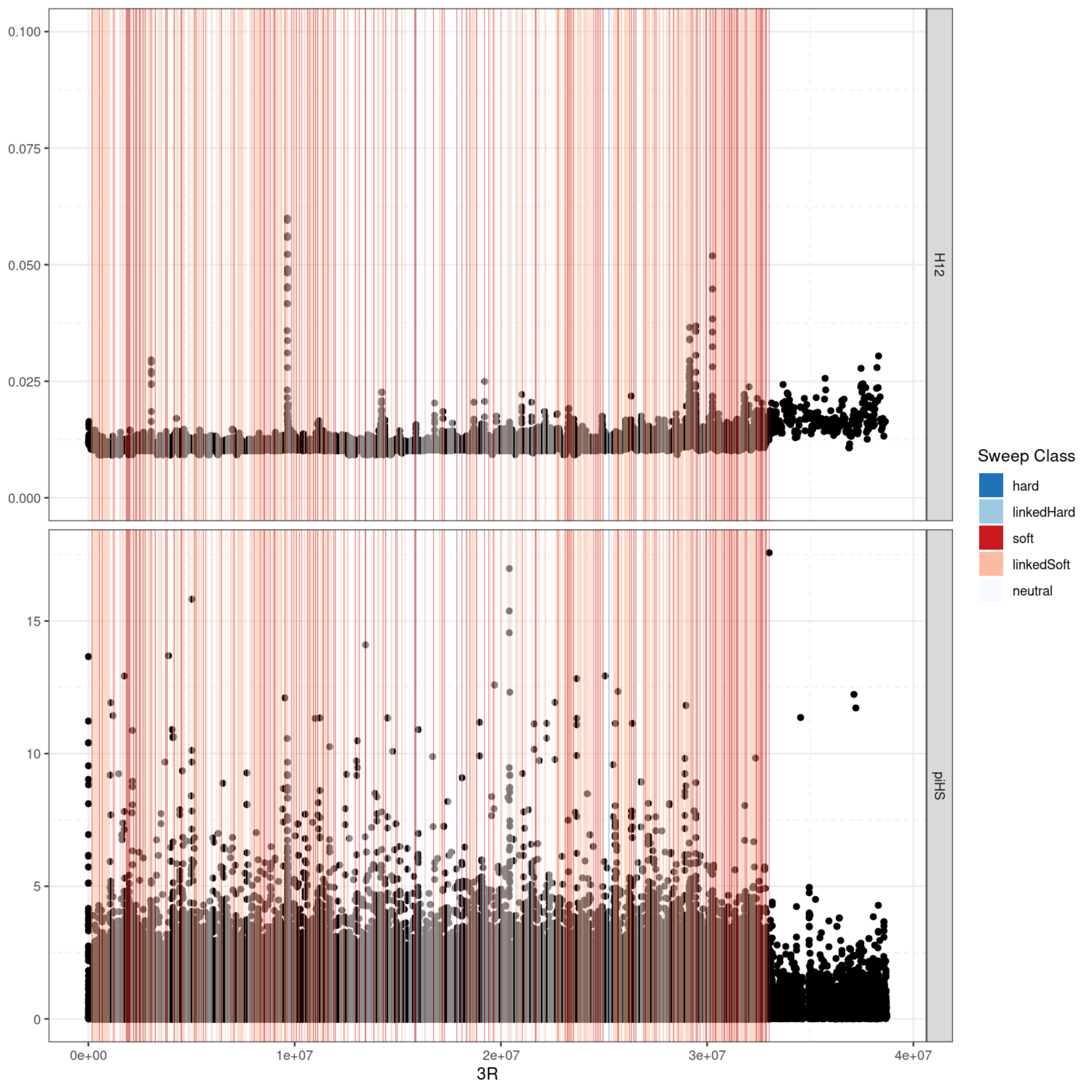

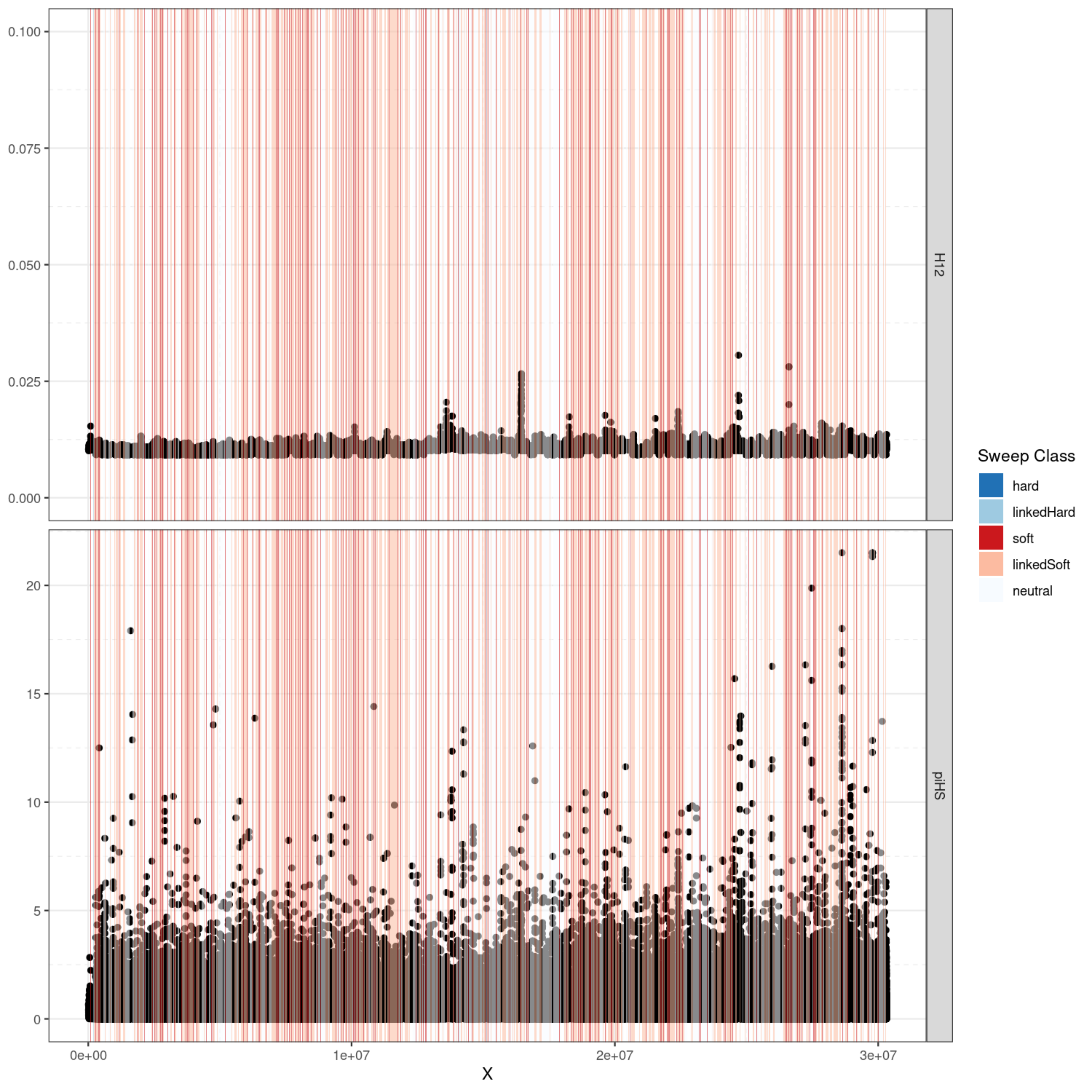

### Supplemental Figure 3

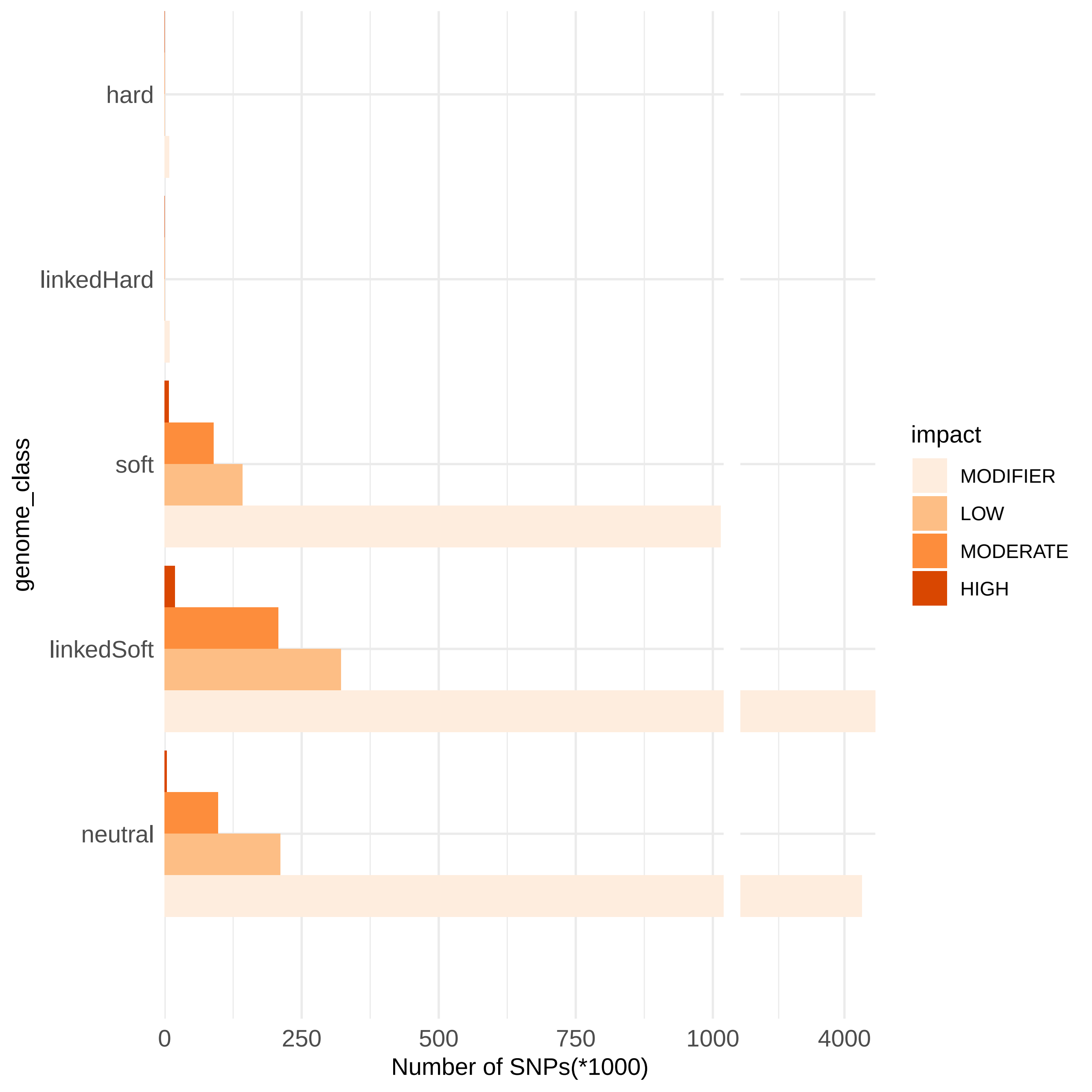
