## Supplemental File 1 for "Direct and indirect impacts of positive selection on genomic variation in *Drosophila serrata*"

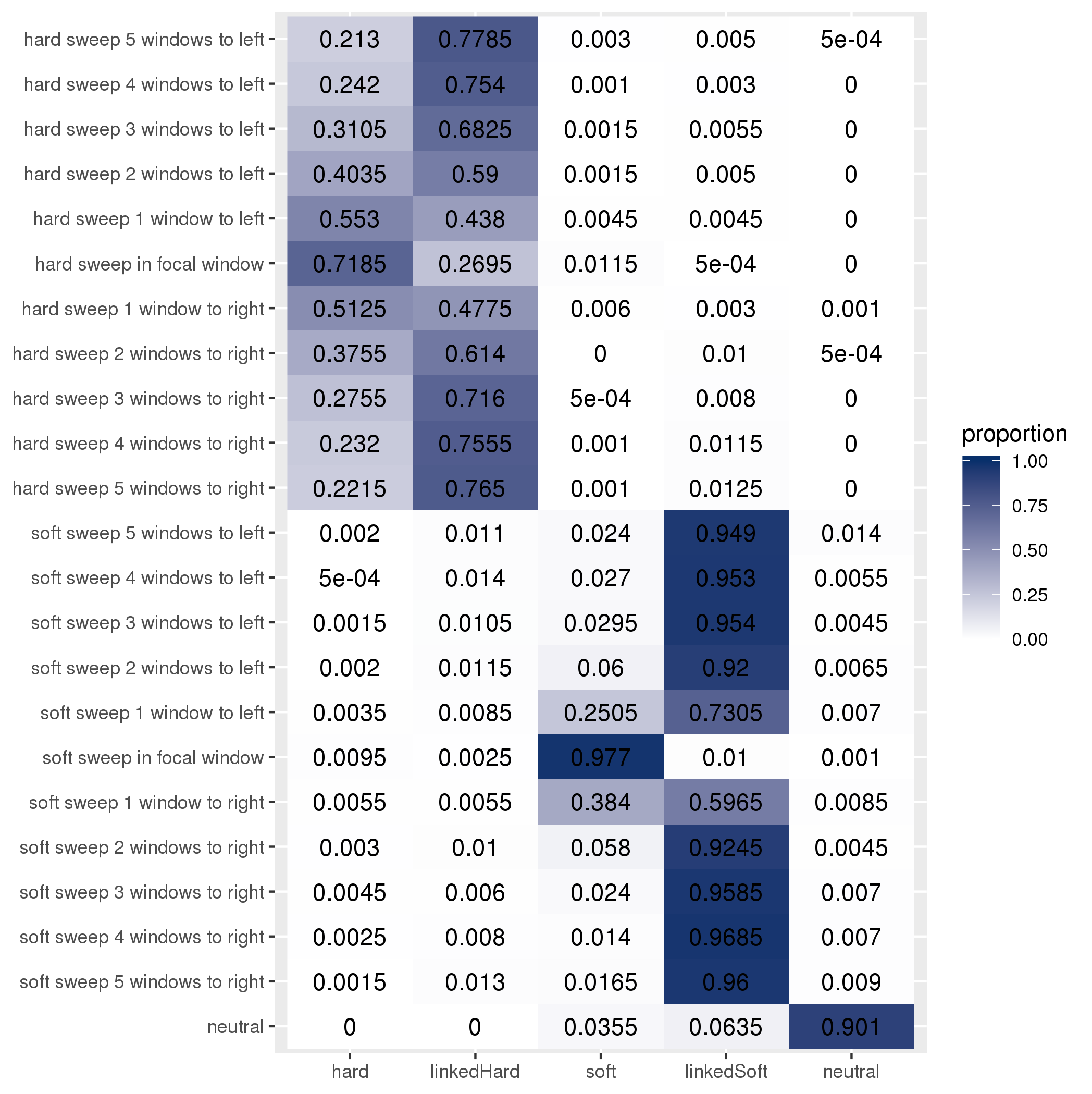

Train: constant population size; test: constant population size

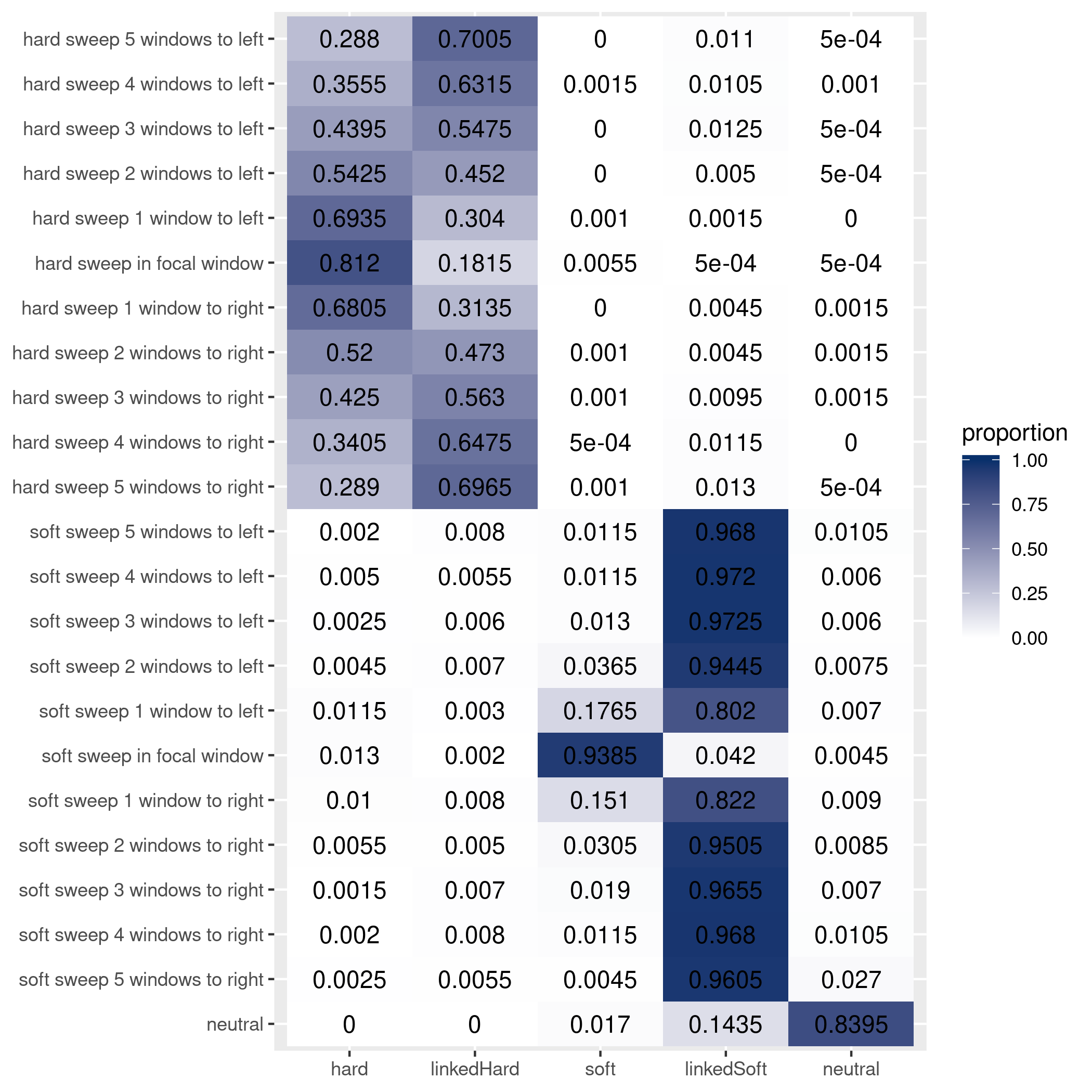

Train: long shallow bottleneck; test: long shallow bottleneck

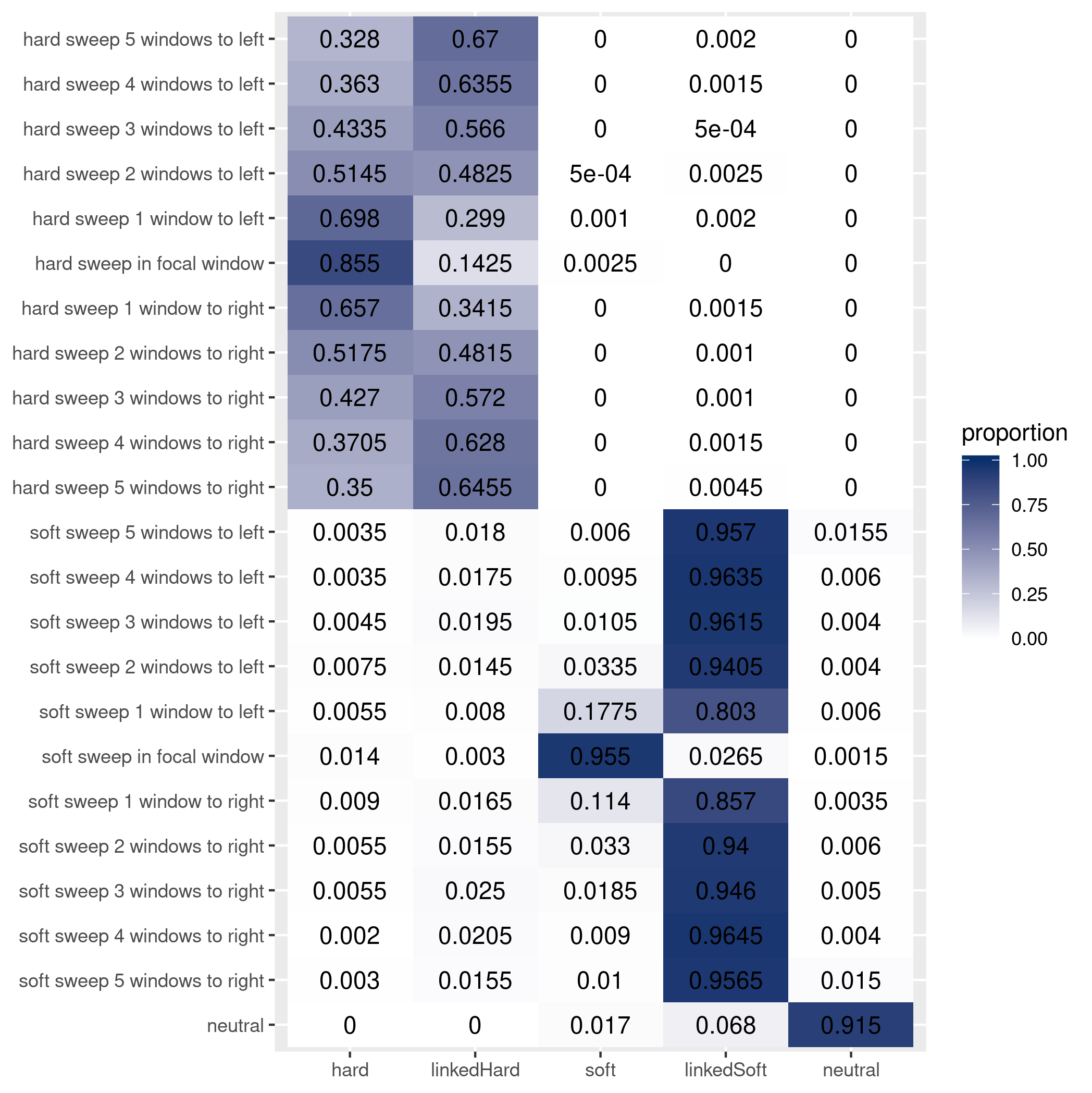

Train: short severe bottleneck; test: short severe bottleneck

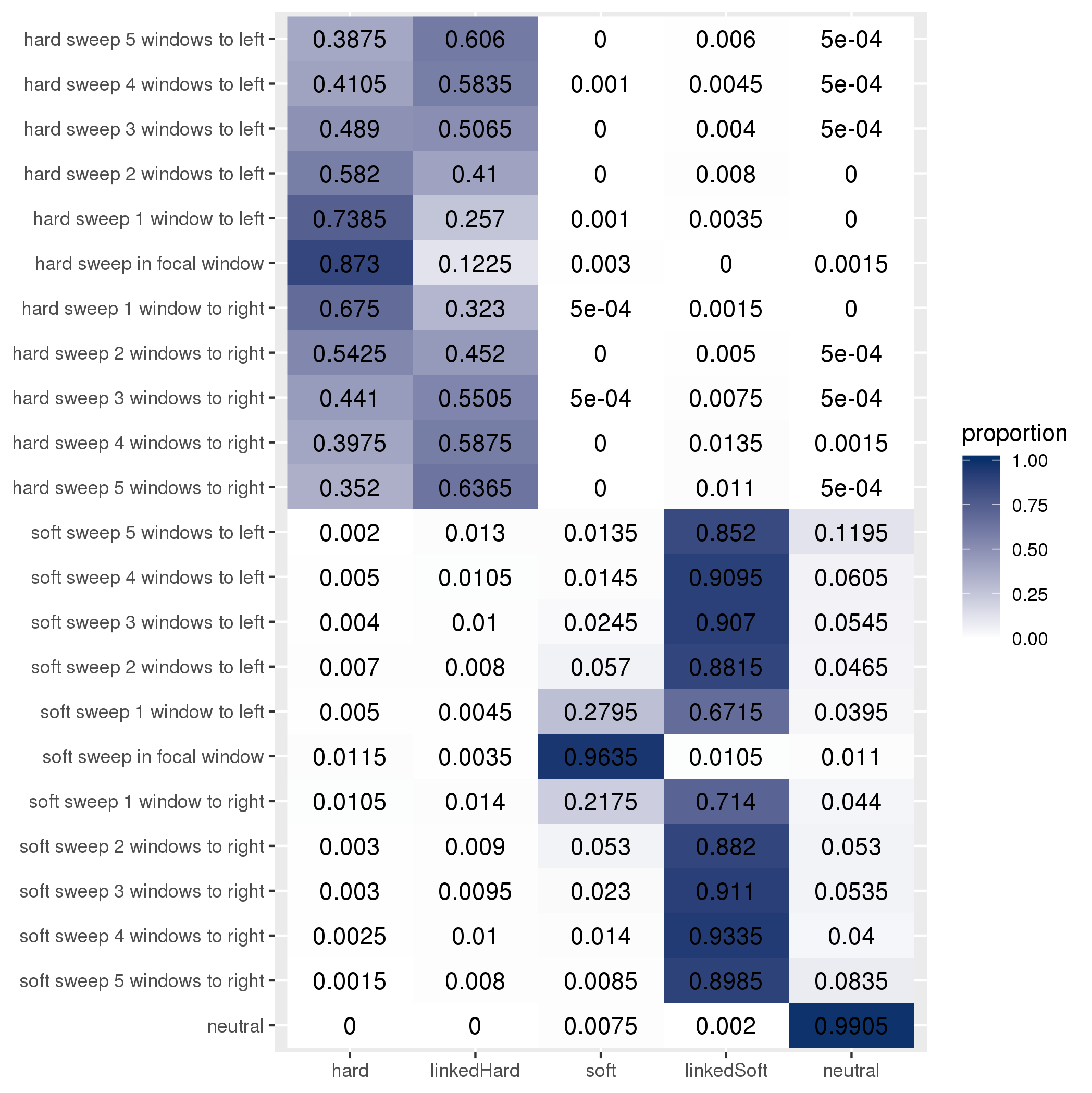

Train: constant population size; test: short severe bottleneck

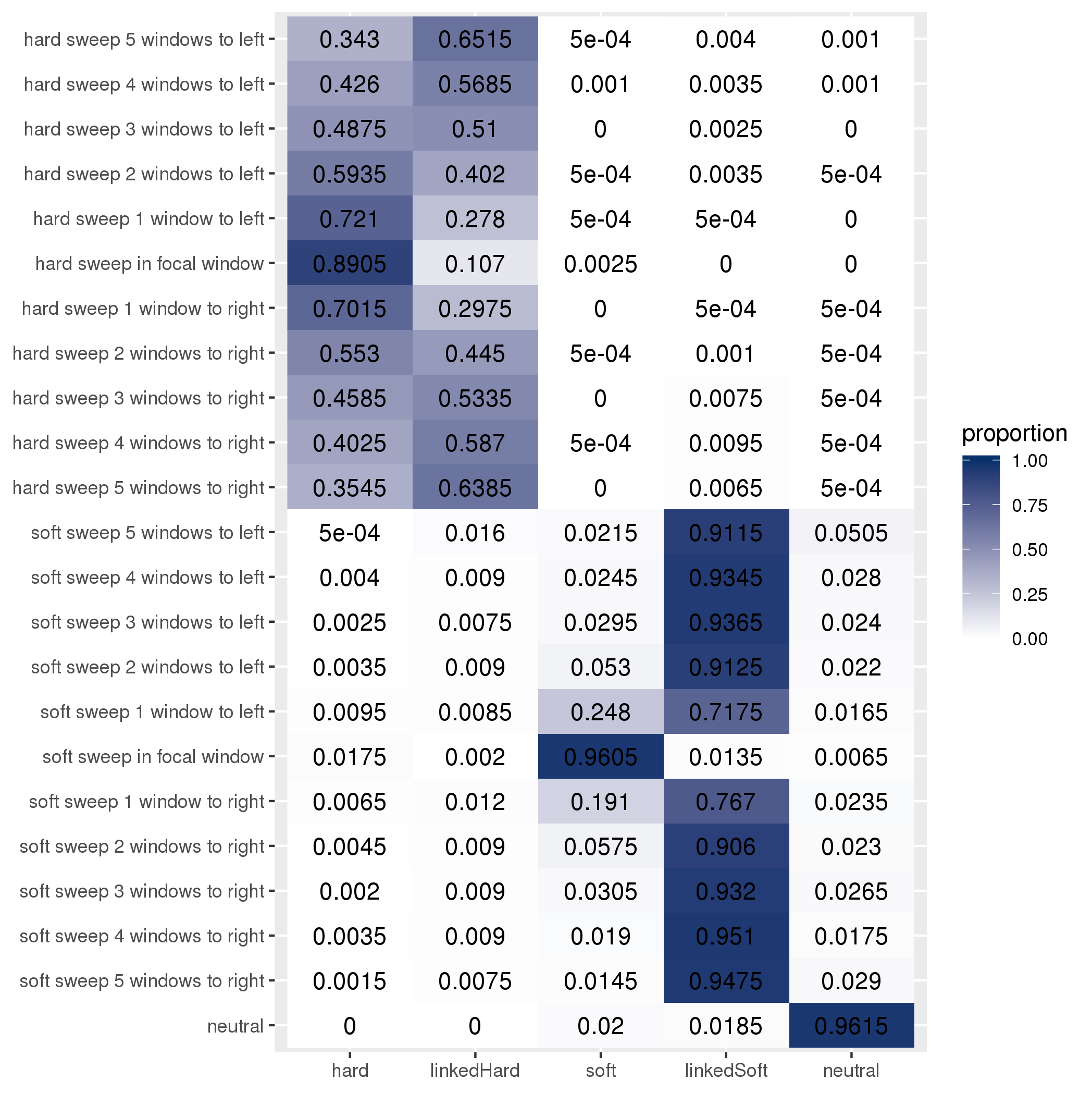

Train: constant population size; test: long shallow bottleneck

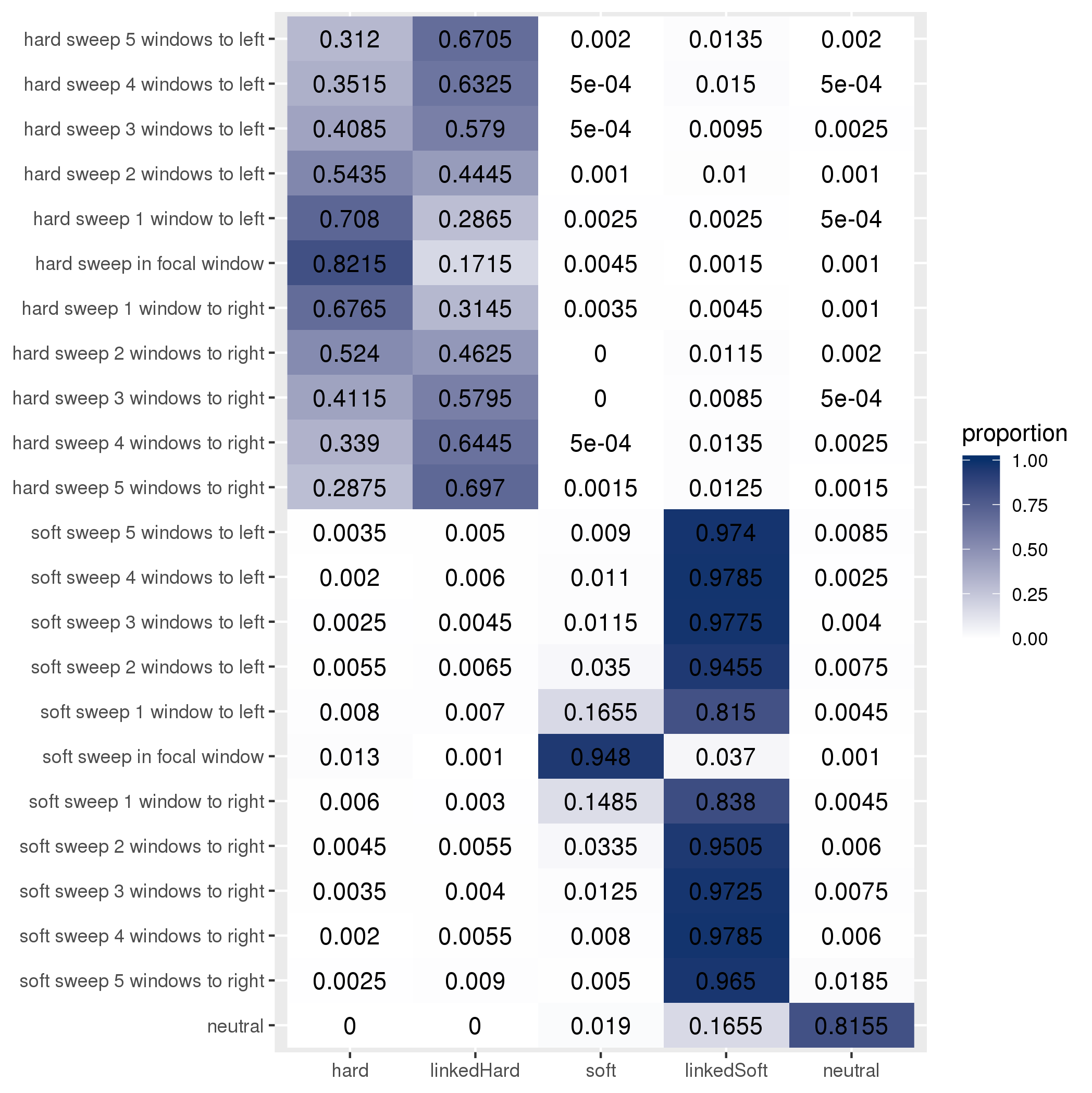

Train: long shallow bottleneck; test: constant population size

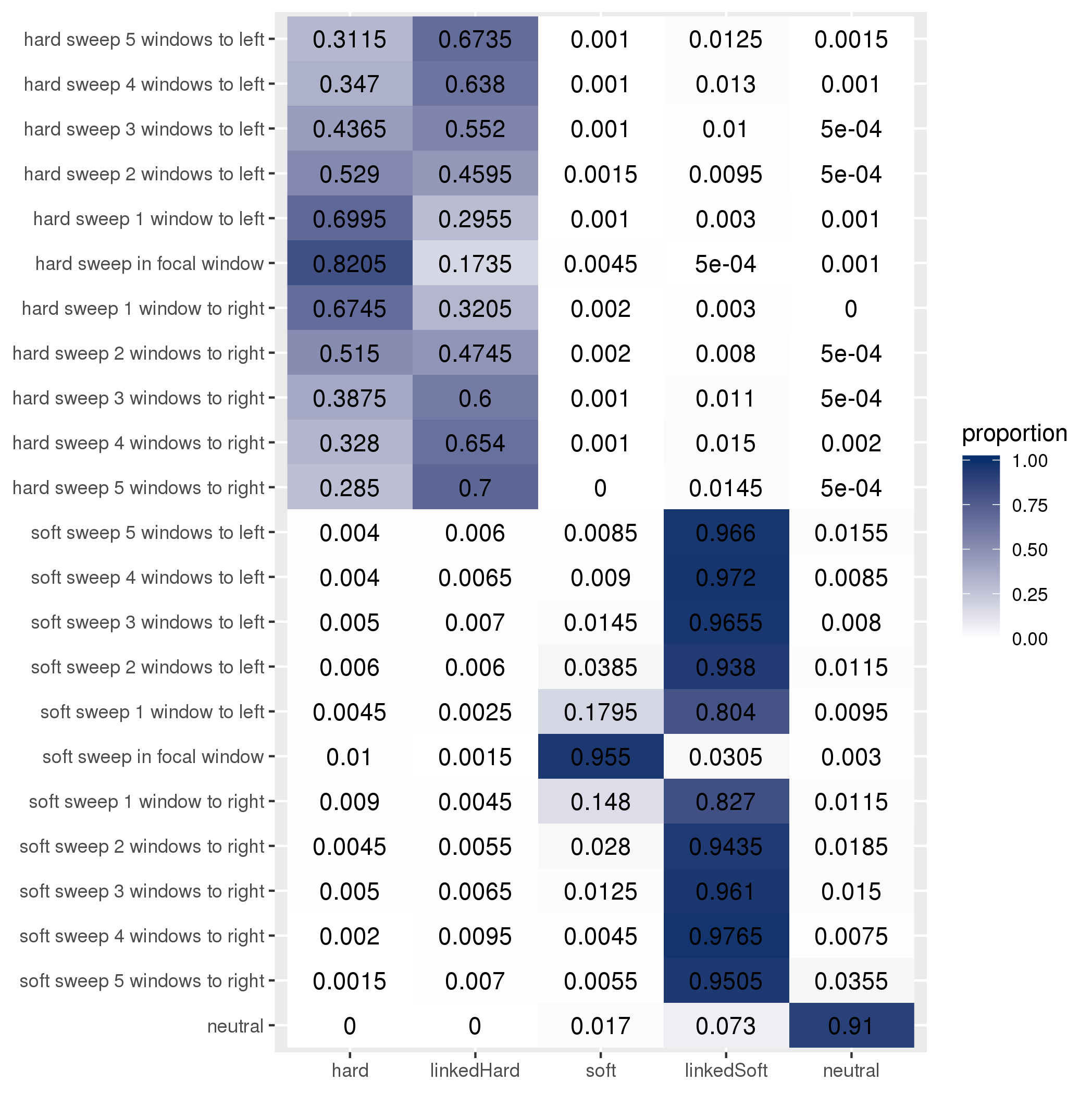

Train: Long shallow bottleneck; test: short severe bottleneck

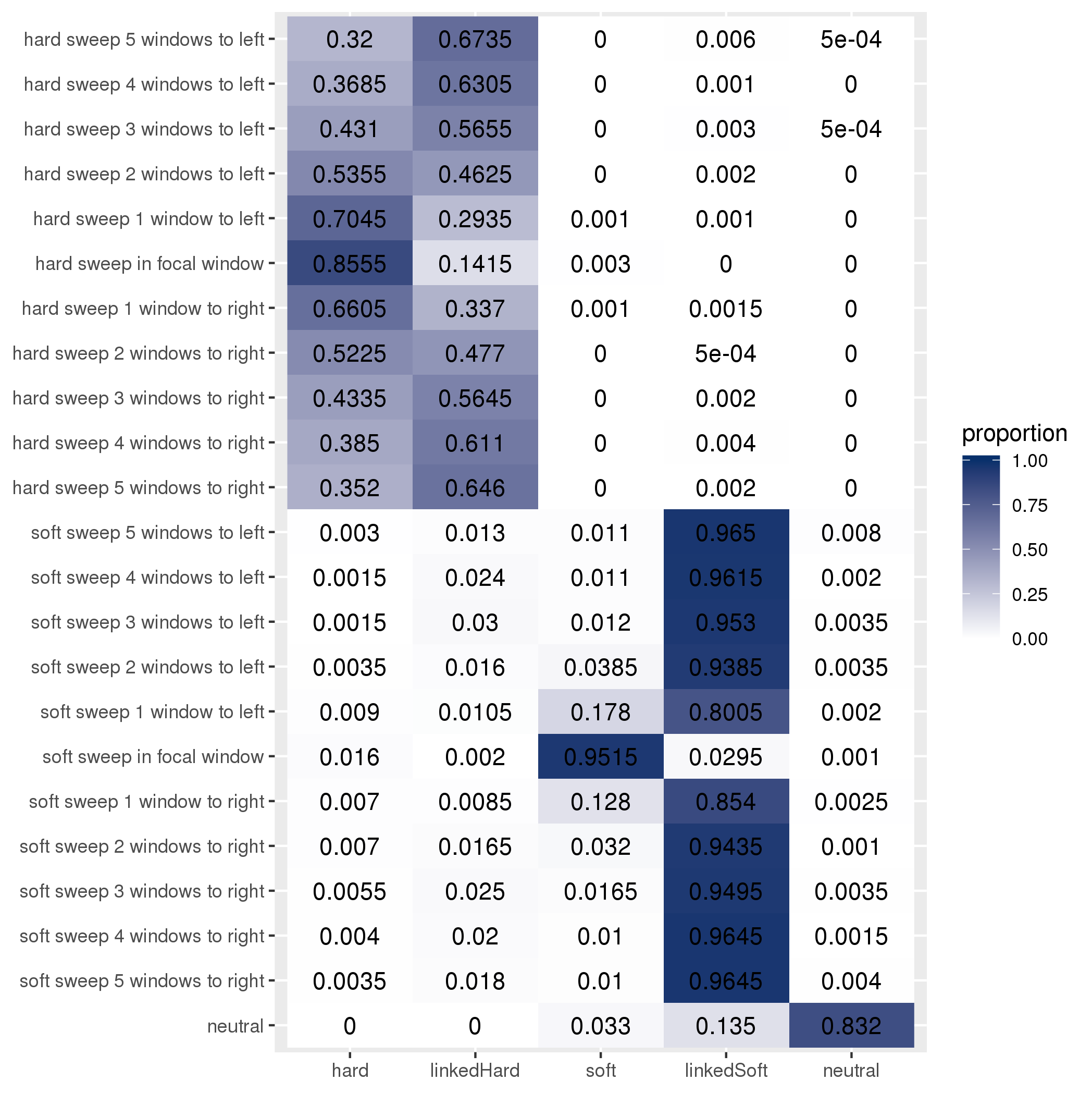

Train: short severe bottleneck; test: constant population size

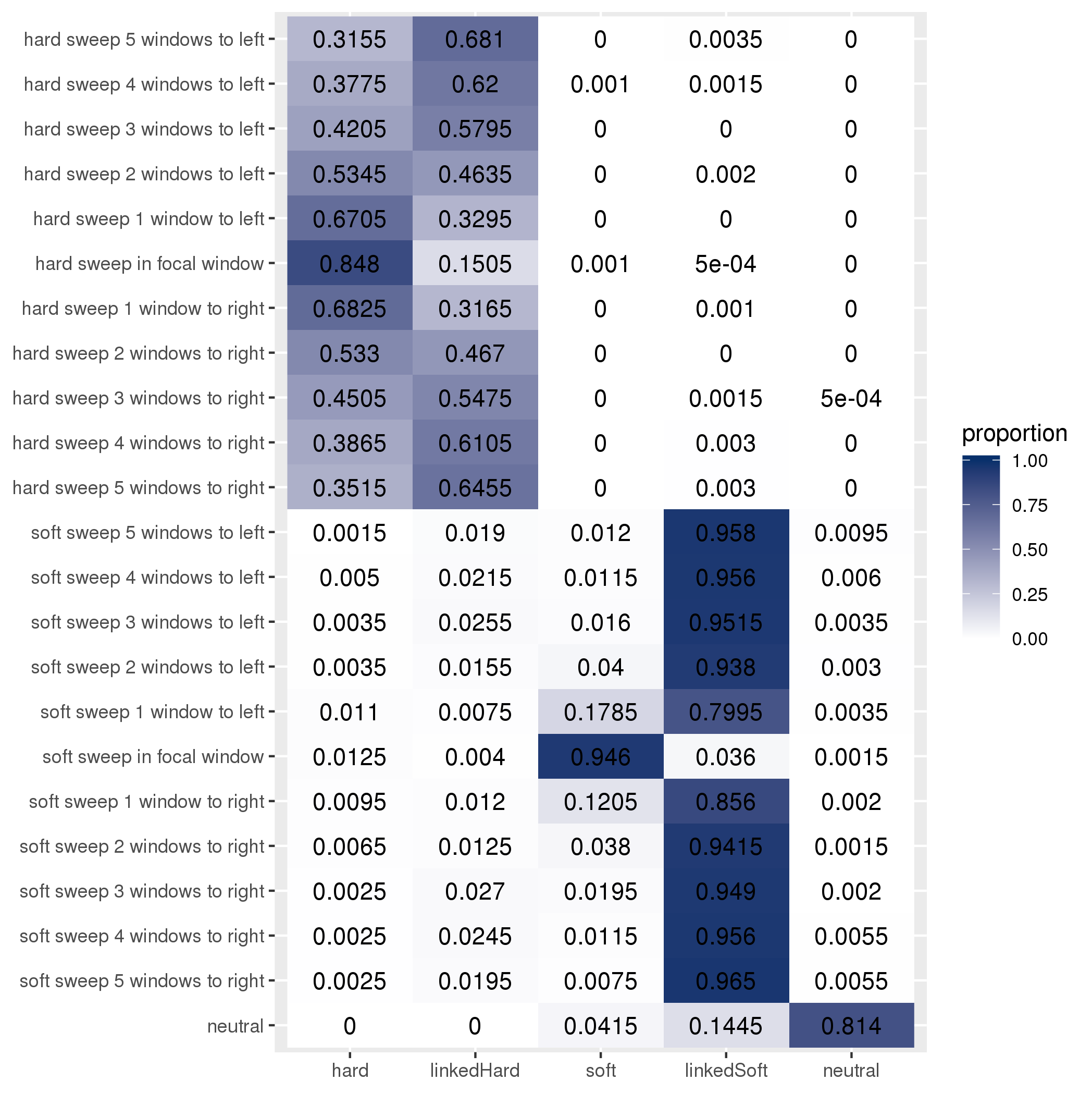

Train: short severe bottleneck; test: long shallow bottleneck
